# Hydrogen Peroxide Production by Epidermal Dual Oxidase 1 Regulates Nociceptive Sensory Signals

**DOI:** 10.1101/2022.09.11.507480

**Authors:** Anna Pató, Kata Bölcskei, Ágnes Donkó, Diána Kaszás, Melinda Boros, Lilla Bodrogi, György Várady, F.S. Veronika Pape, Benoit Roux, Balázs Enyedi, Zsuzsanna Helyes, Fiona M. Watt, Gábor Sirokmány, Miklós Geiszt

## Abstract

Keratinocytes of the mammalian skin provide not only mechanical protection for the tissues, but also transmit mechanical, chemical, and thermal stimuli from the external environment to the sensory nerve terminals. Sensory nerve fibers penetrate the epidermal basement membrane and function in the tight intercellular space among keratinocytes. Here we show that epidermal keratinocytes produce hydrogen peroxide upon the activation of the NADPH oxidase dual oxidase 1 (DUOX1). This enzyme can be activated by increasing cytosolic calcium levels. Using DUOX1 knockout animals as a model system we found an increased sensitivity towards certain noxious stimuli in DUOX1-deficient animals, which is not due to structural changes in the skin as evidenced by detailed immunohistochemical and electron-microscopic analysis of epidermal tissue. We show that DUOX1 is expressed in keratinocytes but not in the neural sensory pathway. The release of hydrogen peroxide by activated DUOX1 alters both the activity of neuronal TRPA1 and redox-sensitive potassium channels expressed in dorsal root ganglia primary sensory neurons. We describe hydrogen peroxide, produced by DUOX1 as a paracrine mediator of nociceptive signal transmission. Our results indicate that a novel, hitherto unknown redox mechanism modulates noxious sensory signals.

## Introduction

Epidermal keratinocytes serve not only as a mechanical barrier to maintain the homeostatic integrity of our body, but they also function as first responders to a broad range of mechanical, chemical, and thermal inputs from the external environment. They possess several membrane receptors and ion channels that trigger intracellular signaling cascades upon activation by various stimuli. Keratinocytes are also described as the source of several paracrine signaling molecules like prostaglandins, cytokines, nucleotides, and also reactive oxygen species (ROS) (Andoh et al., 2001; Ansel et al., 1990; Burrell et al., 2005; Sirokmány et al., 2016). These secreted substances can mediate intercellular communication in the skin.

The detailed histological analysis of mammalian skin in previous studies revealed the presence of unmyelinated nerve fibers in the epidermis. These polymodal sensory nerve fibers penetrate the dermal-epidermal basement membrane and interweave among the very tightly organized epidermal cell layers. Epidermal nerve fiber density (ENFD) is a clinically important histological marker in several neuropathic conditions (Porubcin and Novak, 2020). The close proximity of nerve fibers and keratinocytes renders the latter suitable to function as a secondary sensory cell that can trigger or modulate the activation of the sensory neuron. The narrow, almost virtual intercellular space between epidermal keratinocytes might contribute to high local concentrations of paracrine mediators around the activated epithelial cells.

Both acute inflammatory and persistent inflammatory and neuropathic pain have previously been reported to be enhanced by the presence of ROS (Khalil et al., 1999; Khattab, 2006; Nagy et al., 2010). Although various leukocyte and neuronal sources of ROS production have been identified that mediate these pro-algesic effects of ROS, relatively little is known about the role of ROS produced by keratinocytes. The biological effects of ROS released locally, frequently, and repeatedly by keratinocytes may significantly differ from the effects of ROS from white blood cells and nerve fibers during invasive tissue injury.

ROS can be formed as a byproduct of several metabolic pathways but there is also a group of enzymes – the NADPH oxidase family - devoted specifically to the regulated production of ROS. Members of the NADPH oxidase family typically do not have a ubiquitous, widespread distribution but rather show a specific, unique expression pattern. According to previous reports and publicly available expression databases DUOX1 is expressed in the mammalian skin (Rashid et al., 2019; Sirokmány et al., 2016). Our goal was to characterize the expression and activity of DUOX1 in the mammalian skin in more detail and reveal its potential biological functions.

Here we show that several different signaling pathways that lead to intracellular calcium signals in keratinocytes can induce the release of H_2_O_2_. We provide evidence that the major source of this calcium-dependent peroxide production of keratinocytes is the NADPH oxidase family member dual oxidase one (DUOX1).

We analyzed the noxious behaviors of wild-type and DUOX1 deficient animals following the application of cutaneous nociceptive stimuli and we found increased thermal hyperalgesia in DUOX1 knockout mice.

Our results indicate that DUOX1-derived H_2_O_2_ induces redox-dependent changes in the Kcnq4 (Kv7.4) voltage-gated potassium channel and may also modulate the Transient Receptor Potential Ankyrin Repeat 1 (TRPA1) non-selective cation channel activity.

Based on our *in vivo* and *in vitro* results we propose a new model where the DUOX1-dependent peroxide production of keratinocytes triggers redox changes in cell membrane potassium channel and TRPA1 channel activity in peripheral nerve fibers of dorsal root ganglia. Thereby keratinocyte-derived H_2_O_2_ can alleviate the sensitivity and responsiveness of sensory nerve fibers in a paracrine fashion.

## Results

### Expression and activity of DUOX1 in keratinocytes

We analyzed the expression of NOX/DUOX NADPH oxidases in the mouse skin and mouse primary keratinocytes as well as in the human immortalized keratinocyte cell line (HaCaT) by quantitative reverse transcription PCR. In the mouse tail and paw skin, and also in primary mouse keratinocytes, we found the highest expression levels for DUOX1 and its auxiliary protein DuoxA1 (Figure 1A-C).

**Figure 1.**
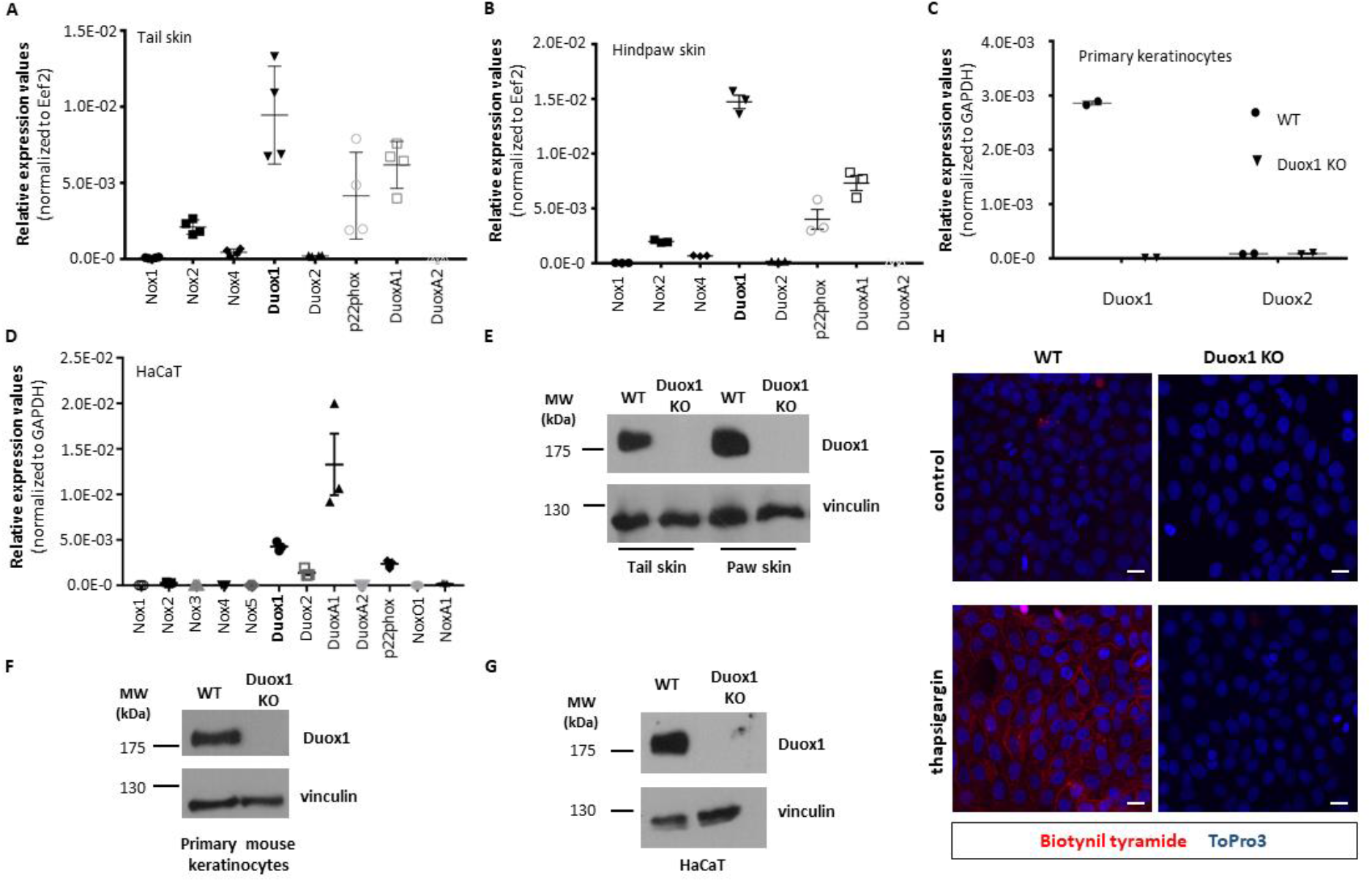
Expression and activity of Duox1 in epithelial cells. Quantitative PCR analysis of the expression of NADPH oxidase family components in mouse tail skin (*A*), hindpaw skin (*B*), primary mouse keratinocytes (*C*) and HaCaT cells (*D*). Dot plots represent mean ± SEM from 3-4 independent experiments. Western blot analysis of Duox1 protein expression in tail skin and hindpaw skin tissue or primary kertinocytes from wild-type and Duox1 knockout mice (*E* and *F*) or HaCaT wild type cells or CRISPR-modified Duox1 knockout cells (*G*). Biotinyl tyramide assay in HaCaT wild-type cells or CRISPR-modified Duox1 knockout cells (*H*). Cells were treated with 1 μM thapsigargin in the presence or absence of horse radish peroxidase. Reaction solution also contained 27.5 μM biotinyl tyramide. After treatment, biotinylated molecules were labeled with fluorescent streptavidin and fixed. Cell nuclei were stained with To-Pro-3. Scale bars: 10 μm.

In accordance with the results of our previous work (Sirokmány et al., 2016) immortalized HaCaT keratinocytes also showed strong DUOX1 expression (Figure 1D). We also analyzed DUOX1 expression at the protein level. As we could not find any commercially available antibody that proved to be specific in DUOX1-knockout controlled experiments, we developed a rabbit polyclonal antibody against a recombinant protein containing amino acids 622-1032 of the human DUOX1 protein. It was expected that the antibody will recognize both the human and mouse samples as the two species show a 91% amino acid identity and 96% similarity in this region of DUOX1. Accordingly, we detected specific DUOX1 western blot signals in mouse skin, mouse primary keratinocytes, and in HaCaT samples as well (Figure 1E-G).

To visualize how H_2_O_2_ is secreted into the intercellular space, we applied a previously described idea (Larios et al., 2001) to devise a novel fluorescent microscope-based method to detect the DUOX1-dependent production of H_2_O_2_ on a cultured monolayer of keratinocytes. We added biotinylated tyramide and horseradish peroxidase (HRP) into the medium of thapsigargin-stimulated or non-stimulated wild-type or DUOX1-deficient HaCaT cells. In this reaction, HRP covalently attaches biotinylated tyramide to tyrosine side chains of cell surface proteins. The reaction takes place in the immediate proximity of H_2_O_2_ production. We then visualized the biotinylated tyramide reagent through incubation with fluorescently labeled streptavidin. Strong tyramide labeling was observed at cell-cell borders of thapsigargin-stimulated wild-type cells (Figure 1H).

We also measured H_2_O_2_ production using the sensitive Amplex Red reagent. To stimulate DUOX1 activity we used reagents that can elicit strong intracellular calcium signals like the endoplasmic reticulum Ca-ATPase inhibitor thapsigargin, the TRPV4 agonist GSK1016790A, and the purinergic receptor agonist ATPγS. We could show that both in mouse primary keratinocytes (Figure 2A) and in HaCaT cells (Figure 2B), genetic deletion of DUOX1 completely eliminated the induced H_2_O_2_ production. These results prove the indispensable role of Duox1 in the ROS production of primary keratinocytes. It is also in agreement with our previous work where we used siRNA knockdown of DUOX1 in HaCaT and A431 cell lines (Sirokmány et al., 2016).

**Figure 2.**
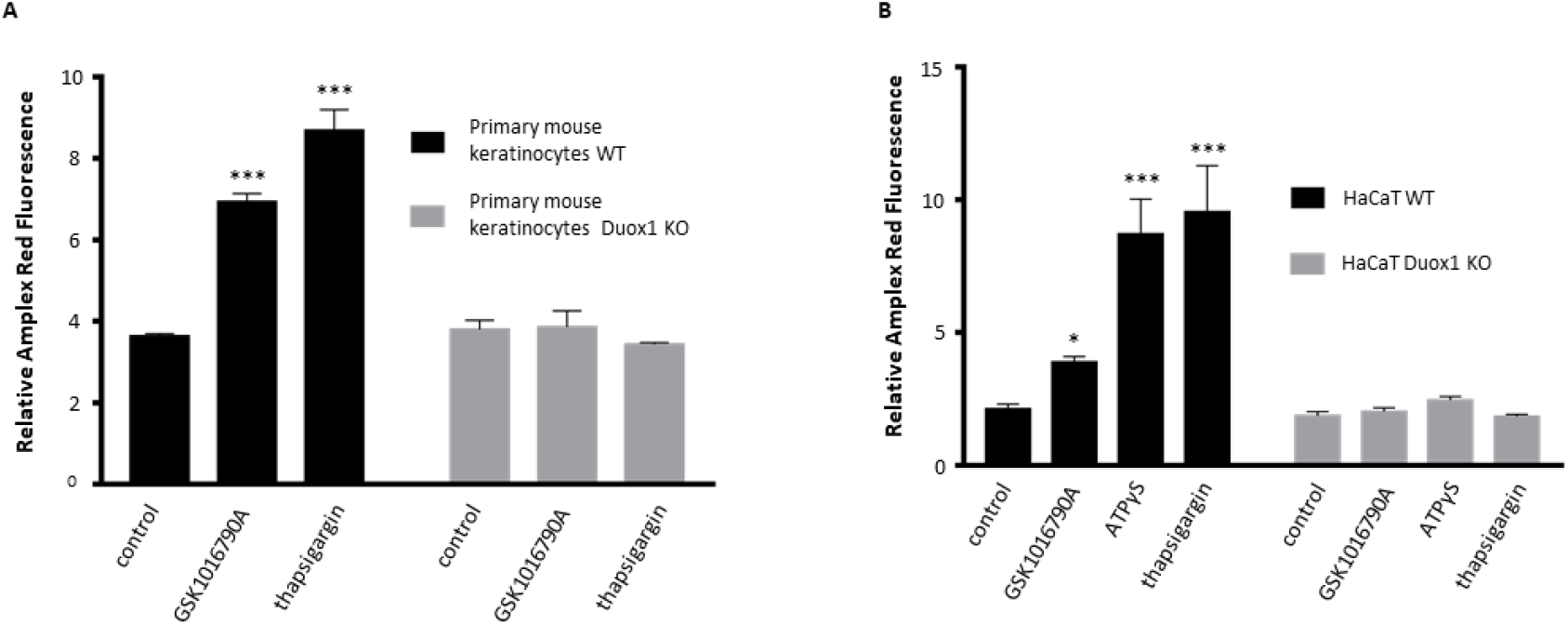
Measurement of H_2_O_2_ production with horseradish peroxidase and Amplex Red reagent. (*A*) Primary mouse back skin keratinocytes were stimulated with 2 nM GSK 1016790A or 100 nM thapsigargin for 30 minutes. (*B*) HaCaT wild-type or CRISPR-modified Duox1 knockout cells were stimulated with 2 nM GSK 1016790A or 10 μM ATPγS or 1 μM thapsigargin for 30 min. Cumulative H_2_O_2_ production after 30 min normalized to initial H_2_O_2_ production. Representative plot (2 independent experiments for primary cells and 3 independent experiments for HaCaT cells) shows mean ± SD of triplicate. * p<0.05, *** p <0.001 compared with corresponding controls, by 2-way ANOVA.

### Histological analysis of DUOX1-deficient mouse skin

To understand if DUOX1 plays a role in physiological sensory functions of the skin we turned to the analysis of DUOX1-deficient animals. As the structural integrity of the skin – or the lack of it – can fundamentally affect sensory signaling processes, we wanted to establish if there is any discernible change in the structure and differentiation of the mouse skin. This analysis was especially vindicated considering the severe cuticle abnormalities described previously in Duox knock-down *C. elegans* (Edens et al., 2001; Kamath et al., 2003). The macroscopic appearance of DUOX1 knockout animals is indistinguishable from wild-type mice. Therefore, we carried out a detailed histological analysis of skin samples of wild-type and DUOX1 knockout mice. Paraffin-embedded tissue sections from mouse tail and paw skin were H&E stained or labeled with specific antibodies (Figure 3A and B). We used a keratin-14 antibody to mark the proliferative basal layer of the stratified epidermis, keratin-10 and loricrin antibodies for early and late differentiation of keratinocytes (Figure 3B) connexin-43 and γ-catenin antibodies to show the intercellular connection between keratinocytes through the gap- and tight junctions respectively (Figure 3 - figure supplement 1). To reveal if there is any possible change in the epidermal organization at the ultrastructural level we studied the stratified epithelium of the mouse tongue by transmission electron microscopic analysis (Figure 3C). Epidermal nerve fibers can be specifically immunolabeled with antibodies against the PGP9.5 (ubiquitin carboxyl-terminal hydrolase L1) antigen. Using 100 μm thick paraffin sections of paw skin epidermis we could visualize the winding intraepidermal nerve fibers. Also, on tail skin wholemount tissue samples, we could immunolabel intracutaneous nerve fibers using anti-Tuj1 antibodies. However, in both immunolabeling techniques, there were no obvious qualitative changes in density or branching of these nerve fibers (Figure 3 - figure supplement 2). Based on all these results we concluded that there was no discernible alteration of epidermal differentiation and structure in the skin of DUOX1 knockout mice.

**Figure 3 with 2 supplements.**
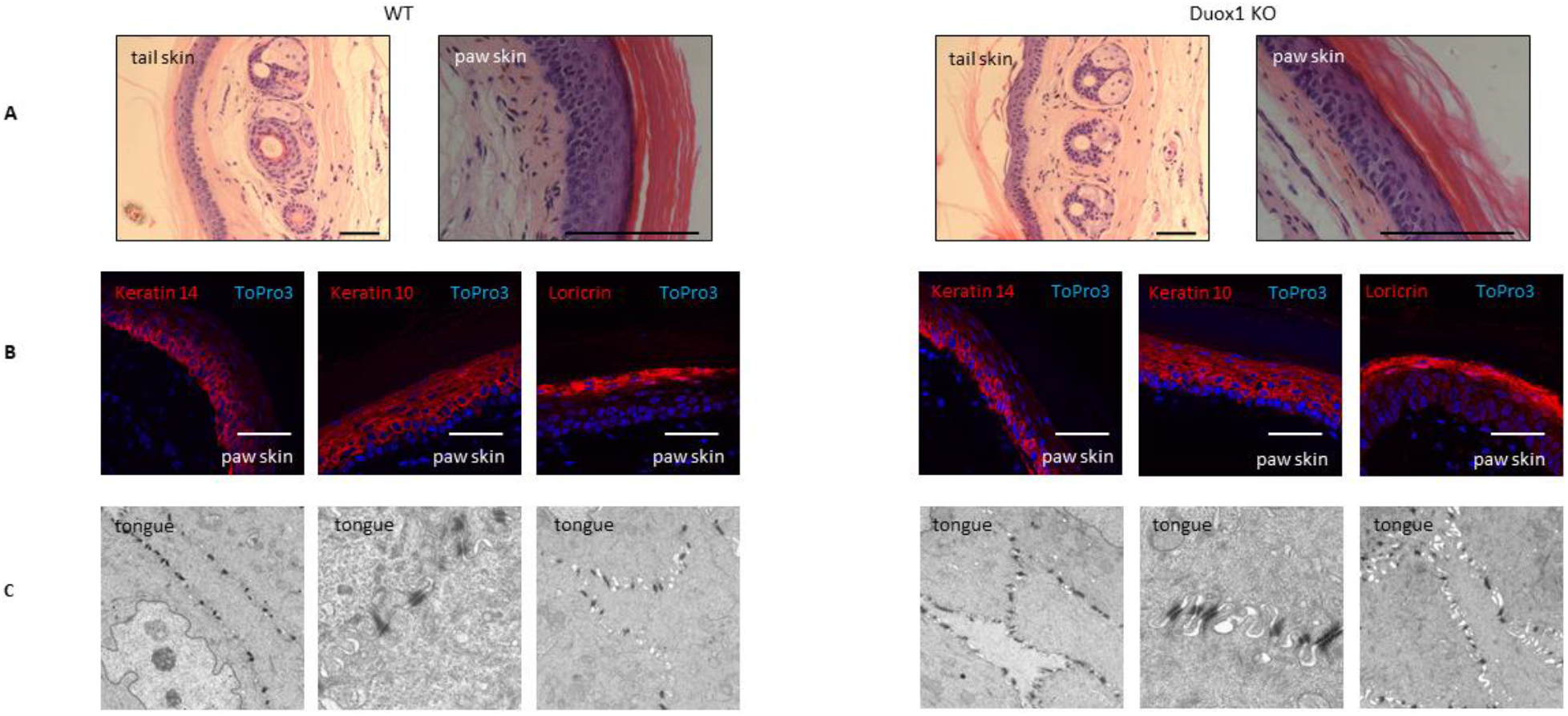
Histological analysis of wild-type and DUOX1-deficient mouse skin. (*A*) H&E staining of paraffin embedded tissue sections from wild-type and Duox1 KO mouse tail and paw skin. Scale bars: 50 μm. (*B*) Wild-type or Duox1 KO mouse paw skin sections were labeled with keratin-14, keratin-10 and loricrin antibodies. Scale bars: 25 μm. (*C*) Analysis of WT and Duox1 KO mouse tongue by transmission electron microscopy.

### Selectively altered nociceptive behavior of DUOX1 deficient animals

As the above results demonstrated that DUOX1 is the predominant source of ROS in keratinocytes we analyzed how DUOX1 activity might affect nociceptive functions in mice. We applied various noxious stimuli to the mouse skin in order to unveil potential differences in the nociceptive behavior of DUOX1 deficient animals.

Pretreatment with the TRPA1 agonist allyl-isothiocyanate (mustard oil) on the mouse skin causes the activation of the capsaicin-sensitive sensory nerve terminals, and release of proinflammatory neuropeptides, inducing acute neurogenic inflammation, and consequently increased sensitivity towards thermal stimuli (Tékus et al., 2016). The decrease of the thermonociceptive threshold (hyperalgesia) is a measurable parameter. As shown in Figure 4, mustard oil pretreatment (5% for 30 sec) of the tail elicited tail removal from the increasing temperature water bath at significantly lower temperatures in DUOX1 knockout animals than in wild-type animals, i.e. the DUOX1 deficient animals were much more sensitized to thermal stimuli than control animals. The difference was visible throughout the entire observation period (10-60 min after the stimuli). This indicated an alleviating effect of DUOX1 activity in the nocifensive behavior elicited by cutaneous thermal stimuli. Importantly, intraperitoneal naloxone pretreatment did not lower the thermal nociceptive threshold in the mice, indicating that endogenous opioid mechanisms do not play a role in this process (Figure 4 – figure supplement 1).

**Figure 4 with 1 supplement.**
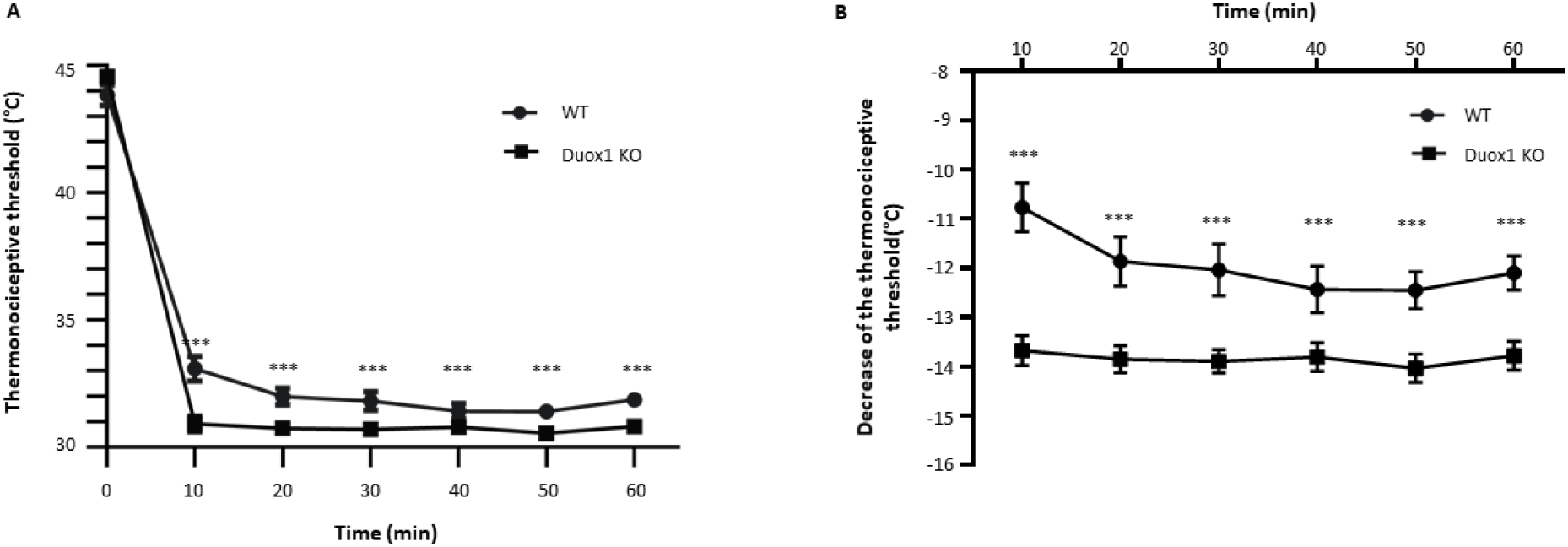
Allyl isothiocyanate-induced thermal hyperalgesia on the tail of wild-type and Duox1 knockout mouse. (*A* and *B*) Thermal hyperalgesia was induced by allyl isothiocyanate (AITC)-contained mustard oil for 30 sec. Comparison of the tail thermonociceptive threshold and its decrease after AITC treatment in wild-type and Duox1 knockout mice. Noxious heat threshold measurements were repeated at 10 min intervals for 60 minutes after treatment. Plots show mean ± SEM of n=16-17 animals/group. *** p <0.001 compared with corresponding controls, by 2-way ANOVA.

We also studied the formalin-induced behavioral response following intraplantar formalin injection. Of note, the nocifensive behavior elicited by formalin treatment also involves TRPA1 activation (Hoffmann et al., 2022). Although we could not observe a significant difference in paw swelling between wild-type and DUOX1 knockout animals (Figure 5 - figure supplement 1), we found an increase in both paw lickings and paw flinches in the second phase of observation (Figure 5A and B) which resulted in a significantly increased composite pain score in DUOX1 deficient animals (Figure 5C).

**Figure 5 with 3 supplements.**
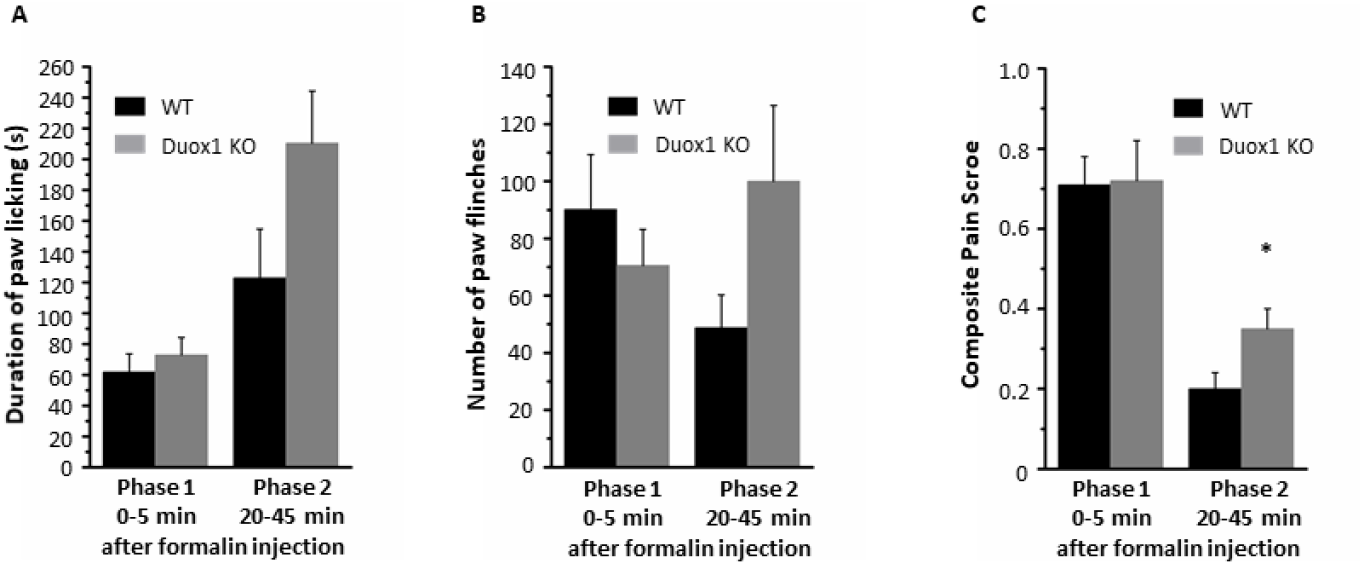
Formalin-induced nociception in wild-type and Duox1 knockout mouse. After intraplantar 20 μl 2.5% formalin injection, spontaneous nocifensive behaviour was observed between 0-5 min and 20-45 min in wild type and Duox1 KO animals. (*A*) The duration of paw licking was measured by a stopwatch. (*B*) Paw flinching responses were counted. (*C*) Composite Pain Score (CPS) was also calculated by the following formula: CPS=(2x paw licking time + 1x paw flinchings)/observation time. Data are mean ± SEM of n=5-5 animals/group. * p<0.05

In the next experiment, neither noxious heat, nor mechanical nociceptive thresholds following subjection of the animals to a short heat injury (immersing the hind paw into a 51 °C water bath for 15 seconds under ether anesthesia) showed significant differences between wild-type and DUOX1 deficient animals (Figure 5 - figure supplement 2).

Finally, following mechanical noxious stimuli (plantar incision) we measured mechanical nociceptive thresholds in both wild-type and DUOX1 knockout groups, but this postoperative hyperalgesia was not different either in the DUOX1 KO animals compared to wild-type controls (Figure 5 - figure supplement 3).

In the following experiments, we aimed to identify the possible molecular targets of DUOX1-mediated H_2_O_2_ production.

### PGE2 and ATP release from stimulated keratinocytes

Keratinocytes can release several mediators that are known to influence nociceptive behavior. We assumed that stimuli that cause an increase in [Ca^++^]_ic_ will simultaneously activate DUOX1 and trigger the secretion of mediators as well. Therefore, in stimulated keratinocytes, we can observe if DUOX1 activity can influence the release of mediators. Intraplantar PGE2 is a known inducer of nociceptive behavior (Keppel Hesselink et al., 2017)and is also described as a secretory product of keratinocytes (Huang et al., 2008). We measured the stable adenosine triphosphate analog ATPγS-elicited PGE2-release of scrambled or DUOX1-specific siRNA treated HaCaT cells. As seen in Figure 6 - figure supplement 1, we could strongly induce the release of PGE2 with ATPγS, however, there was no difference between scrambled and DUOX1 knockdown cells that could explain the previously described behavioral phenotype changes.

**Figure 6 with 4 supplements.**
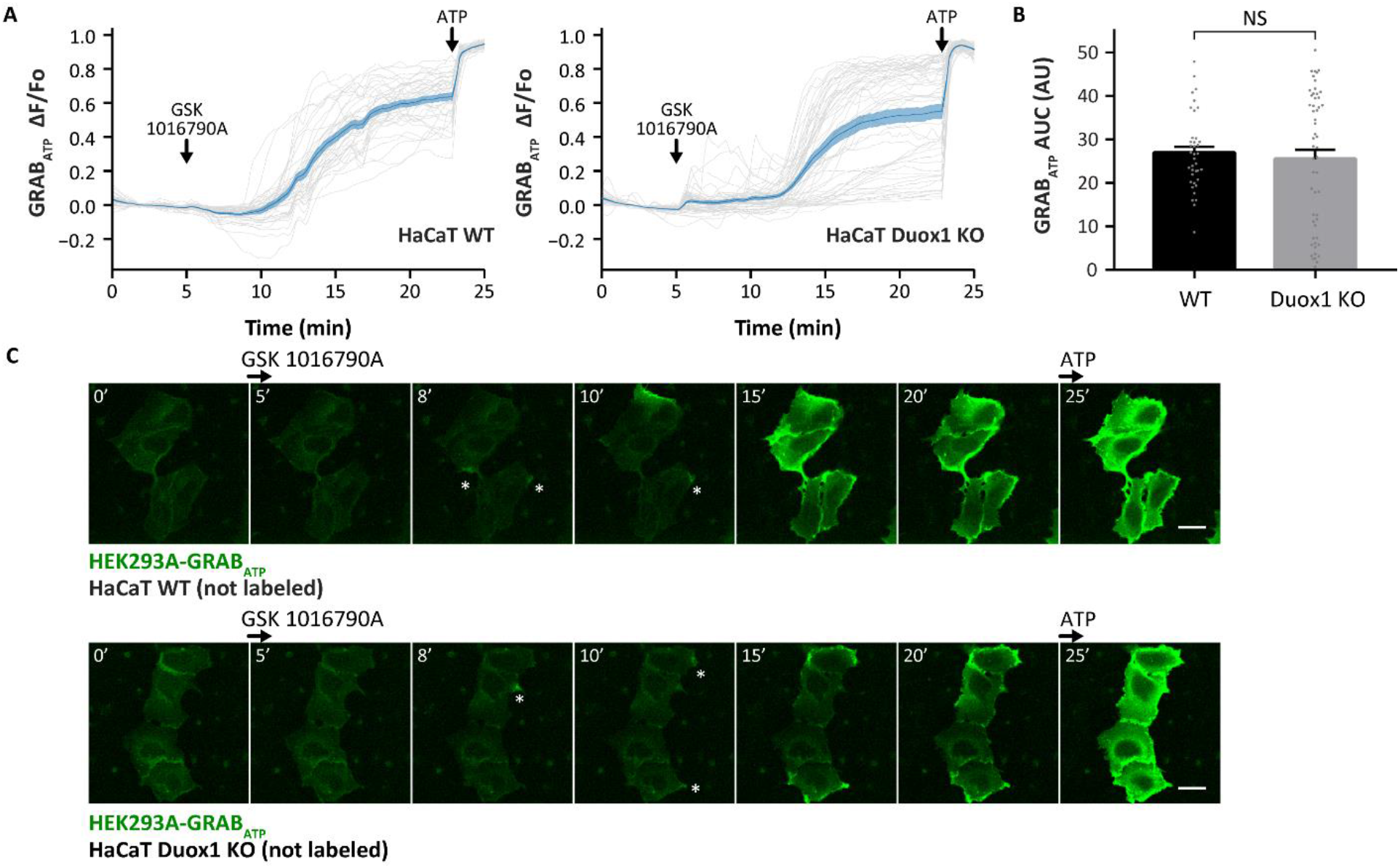
TRPV4 agonist, GSK 1016790A induced ATP secretion from wild-type and Duox1 knock-out HaCaT cells. (A) Extracellular ATP-sensitive, fluorescent GRABATP sensor expressing HEK293A cells were cocultured with WT or Duox1 knock-out HaCaT. After a wash to remove any residual ATP in the media, the cells were treated with 2.5 nM GSK 1016790A and then with 1 μM ATP, as a positive control. Gray lines represent the normalized mean GRABATP intensity of every cell over time. Blue line shows the overall average of the normalized mean intensity of all the cells ± SEM (WT: n = 38, KO: n = 53 cells from 3 independent experiments). (B) The area under the curve (AUC) was measured for each cell during the time of the GSK 1016790A stimulus, between 5-27 min and was plotted per condition (mean ± SEM, gray dots represent values of individual cells). (C) Fluorescent images from different time points showing local oscillations (marked by asterisks) and sustained elevations of the GRABATP signal. Scale bars: 25 μm.

ATP is also a frequently studied peripheral nociceptive mediator (Caterina and Julius, 1999) that can be released from keratinocytes (Mizumoto et al., 2003). However, despite our repeated efforts, with the use of commercially available, luminescence-based ATP-release kits we could not measure significant and reproducible ATP signals in the supernatant of keratinocytes. Therefore, we applied a previously described, genetically encoded, GPCR activation-based ATP sensor, GRAB_ATP_(Wu et al., 2022). This sensor construct is based on the insertion of a circularly permutated enhanced GFP (cpEGFP) into the ATP-binding hP2Y_1_ receptor. The binding of ATP by this fusion construct specifically and sensitively enhances the fluorescence of cpEGFP. We set up a coculture system, where wild-type or DUOX1 deficient HaCaT cells were cocultured with GRAB_ATP_ expressing HEK293A cells. In this system, we selectively stimulated the keratinocytes with the TRPV4 agonist GSK1016790A. This compound does not act on HEK293A cells but induces a strong increase in [Ca^++^]_ic_ and DUOX1 activity in keratinocytes (Luo et al., 2018; Sirokmány et al., 2016). Accordingly, GSK1016790A did not elicit any changes in the fluorescence of GRAB_ATP_ expressing HEK293A cells. H_2_O_2_ treatment did not affect the fluorescence of GRAB_ATP_ either (Figure 6 – figure supplement 2 and Figure 6 - figure supplement 4 video). Thus we assumed that if GSK1016790A triggers ATP-release then it should specifically activate the GRAB_ATP_ sensors on the membrane of the transfected HEK293A cells. Indeed, we could repeatedly observe transient local oscillations followed by sustained high signals of extracellular ATP in the GRAB_ATP_ expressing HEK293A cells following TRPV4 stimulation of the neighboring keratinocytes (Figure 6 and Figure 6 – figure supplement 3 video). However, there were no obvious differences between sensor cell activation cocultured with wild-type or DUOX1-deficient HaCaT cells (Figure 6).

### Hydrogen peroxide-mediated redox changes of TRPA1-dependent intracellular calcium signals

Dorsal root ganglion (DRG) neurons express numerous ion channels and membrane receptors that regulate their excitability. Detailed quantitative RNA-sequencing datasets are available about the membrane receptors and ion channels expressed in these neurons (Manteniotis et al., 2013). We confirmed the strong expression of several of these ion channels in our own qPCR measurements. Importantly, we could not detect any significant changes in the expression level of these genes between wild-type and DUOX1 KO DRG samples (Figure 7A). We also analyzed the expression of TRPV3 and TRPV4 in wild-type and DUOX1 knockout mouse skin samples as these heat sensitive receptors have been previously described to play a role in the thermosensory function of the skin (Huang et al., 2011a; Wetsel, 2011). However, as shown in Figure 7B we did not see a DUOX1-dependent change in the expression of these genes. In the following experiments, we aimed to identify molecular targets, the activity of which might be affected by hydrogen peroxide.

**Figure 7.**
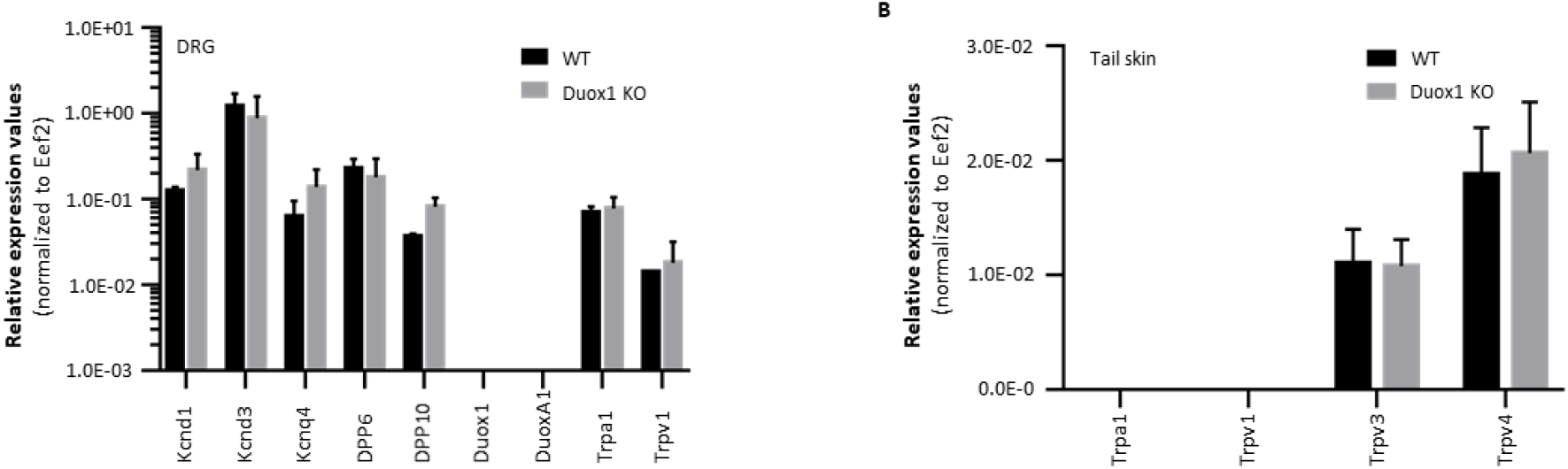
Expression of ion channels and Duox1 in dorsal root ganglion and tail skin. (*A*) Quantitative PCR analysis of the expression of potassium channels, Duox1, DuoxA1 and TRP channels in wild-type and Duox1 knockout mouse dorsal root ganglion. Bars represent mean ± SEM from 2 independent experiments. (*B*) Quantitative PCR analysis of the expression of TRP channels in mouse tail skin. Bars represent mean ± SEM from 2-4 independent experiments.

We browsed the available literature data to find cell surface proteins of DRG cells that had been reported to show redox-sensitive activity. The H_2_O_2_ -sensitive TRPA1 channel is present in certain populations of DRG cells(Andersson et al., 2008).

Therefore, first we expressed recombinant TRPA1 channels in HEK293T cells, and followed intracellular calcium signals upon treatment with H_2_O_2_ and a TRPA1 agonist. As seen in Figure 8A and Figure 8 – figure supplement 1A, the TRPA1-expressing HEK cells - in contrast to the non-transfected cells - showed a continuous, H_2_O_2_-dependent increase in [Ca^++^]_ic_. Importantly, following preincubation with H_2_O_2_, the TRPA1-expressing cells responded to the TRPA1 agonist allyl isothiocyanate (AITC) with a much smaller relative increase in [Ca^++^]_ic_ than those cells without an H_2_O_2_ pretreatment (Figure 8B and Figure 8 – figure supplement 1B). This is a clear indication that H_2_O_2_ has a significant impact on the responsiveness of TRPA1-expressing cells to specific stimuli. These data confirm the idea of keratinocyte-derived H_2_O_2_ acting as a paracrine mediator on TRPA1 expressing sensory nerve fibers.

**Figure 8 with 1 supplement.**
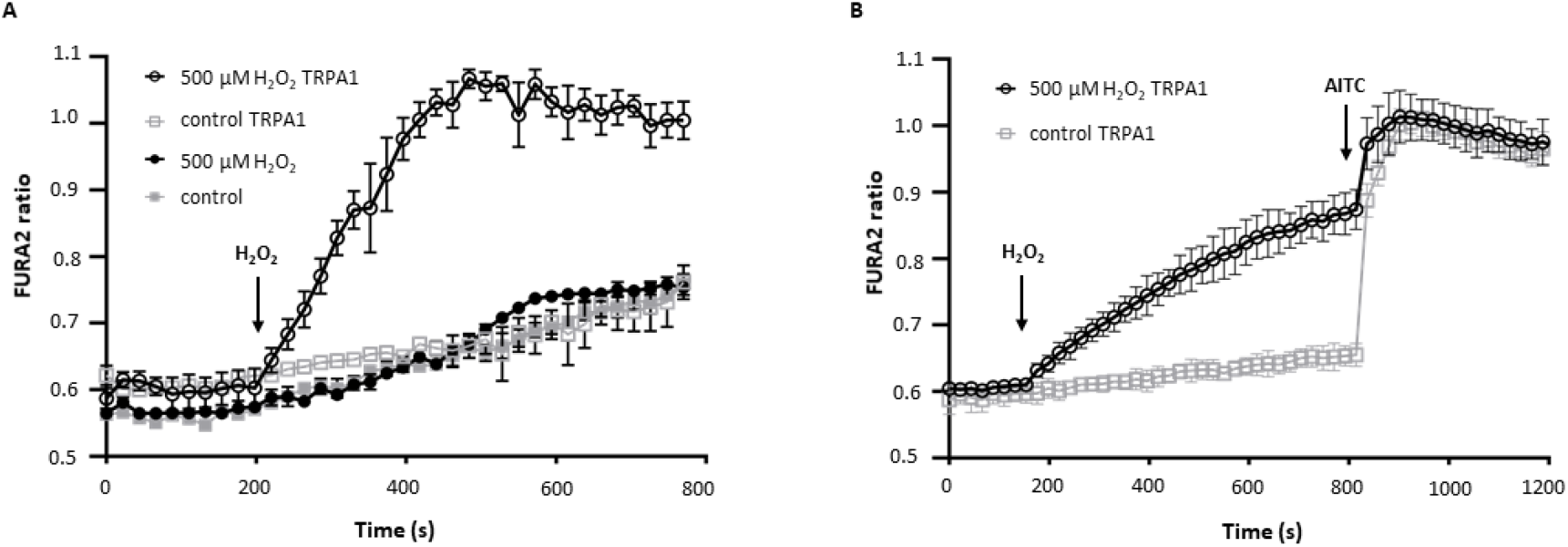
Ca^2+^ measurements with Fura-2-AM in HEK293 cells expressing TRPA1. (*A*) TRPA1-dependent effect of H_2_O_2_ on [Ca^2+^]_ic_. [Ca^2+^]_Ic_ responses evoked by 500 μM H_2_O_2_ in the presence of TRPA1. (*B*) Change in [Ca^2+^]_ic_ of TRPA1 transfected HEK cells in response to sequential applications of 500 μM H_2_O_2_ and 10 μM AITC. Representative plot from at least 3 independent experiments shows the mean ± SD of triplicate.

### Expression and H_2_O_2_ mediated redox changes of Kcnq4 M-type potassium channel

Our next candidate of redox-sensitive ion channels was the voltage-gated M-type potassium channel – Kv7.4 or Kcnq4 – as it was reported in a detailed electrophysiological study to remain in its open state longer after H_2_O_2_-mediated oxidation (Gamper et al., 2006). Our qPCR analysis showed that this channel is expressed – in sharp contrast to DUOX1 – very strongly in primary sensory neurons but was hardly detectable in paw and tail skin samples (Figure 9A and B). We aimed to provide biochemical evidence about the H_2_O_2_-mediated oxidation of cysteine-thiols in this channel protein. We used biotinylated iodoacetamide (BIAM) to follow the oxidation of the FLAG-tagged, recombinant Kcnq4 potassium channel (Brewer et al., 2015). In the first set of experiments, we treated Kcnq4-FLAG transfected HEK293 cells with increasing concentrations of hydrogen peroxide and lysed them in the presence of BIAM. Then we carried out an immunoprecipitation with monoclonal anti-FLAG antibody and ran the immunoprecipitate on SDS polyacrylamide gel to test the amount of biotin labeling with streptavidin-HRP. As it is shown in Figure 9C, the biotin signal decreased as the added H_2_O_2_ concentration increased indicating the oxidation of sulfhydryl groups of Kcnq4. To confirm these results, and investigate the reversibility of oxidation, we carried out BIAM labeling also in a reverse fashion (Löwe et al., 2019). In this case, we first alkylated all the non-oxidized sulfhydryl groups after the peroxide treatment using N-ethyl-maleimide (NEM). Subsequently, we reduced all the reversibly oxidized sulfhydryl groups (aka sulphenic acid) with dithiothreitol and carried out the BIAM labeling of reduced cell lysate samples. Then we continued with anti-FLAG immunoprecipitation and streptavidin-HRP detection as described above. This way the streptavidin-HRP signal increased proportionally with the increased oxidation of the potassium channel (Figure 9D).

**Figure 9.**
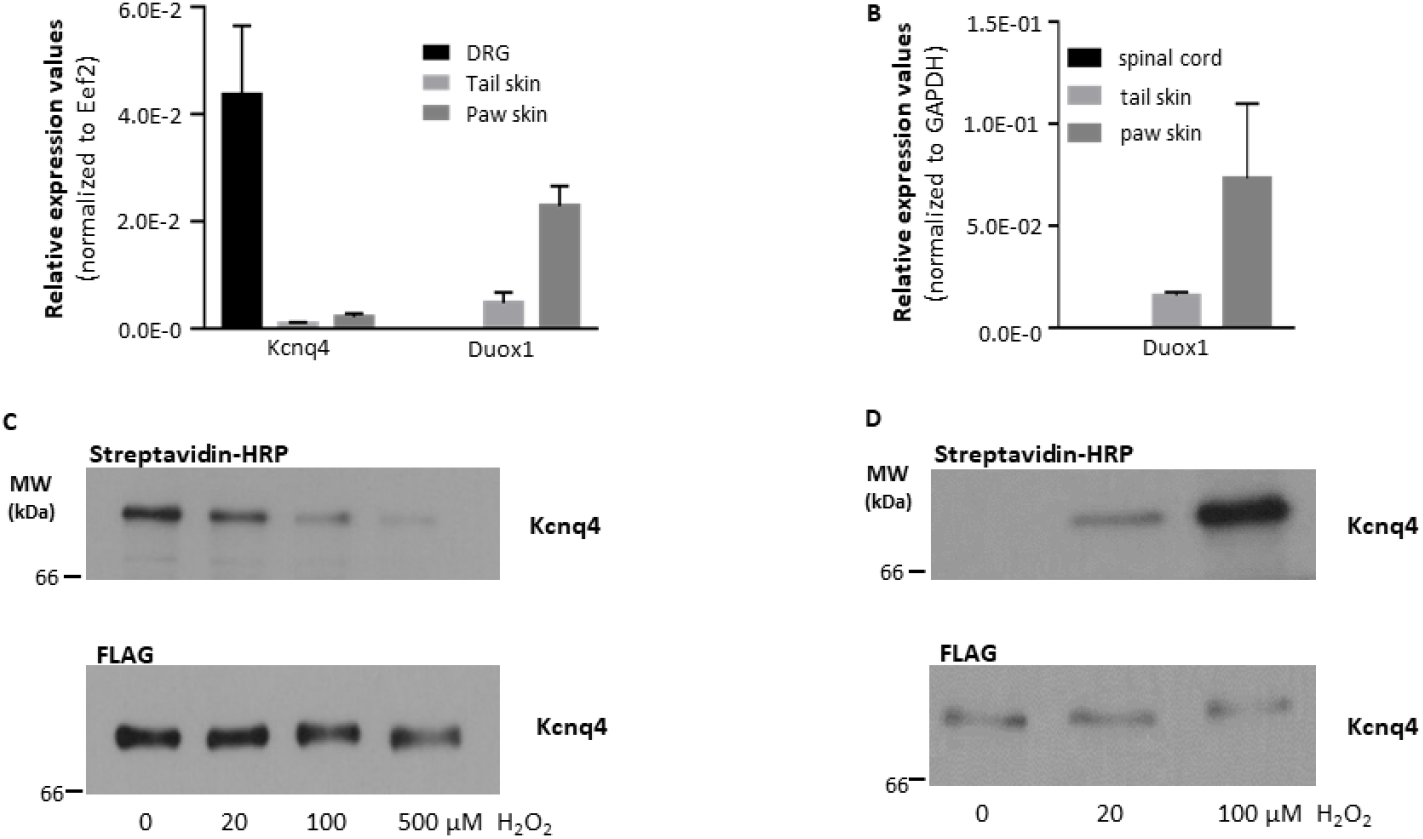
Hydrogen peroxide mediated redox changes of voltage-gated potassium channel, Kcnq4. (*A*) Quantitative PCR analysis of the Kcnq4 and Duox1 in wild-type mouse dorsal root ganglion, tail skin and paw skin. Bars represent mean ± SEM from 3 independent experiments. (*B*) Expression of Duox1 in spinal cord, tail and paw skin. (*C*) HEK293 cells expressing FLAG-tagged Kcnq4 were treated with H_2_O_2_ (0, 20, 100, 500 μM) and lysed in presence of BIAM. Following anti-FLAG immunoprecipitation, BIAM signal was detected by streptavidin-HRP on western blot. (*D*) After H_2_O_2_ treatment (0, 20, 100 μM) non-oxidized thiols were alkylated with N-ethylmaleimide, then lysates were reduced with dithiothreitol and labelled with BIAM. We continued with anti-FLAG immunoprecipitation and streptavidin-HRP detection.

In the later version of redox labeling, we observed that high peroxide concentrations (500 μM) eliminated the streptavidin signal indicating an irreversible oxidation – formation of sulfinic or sulfonic acid – of the thiol groups of Kcnq4 which were impossible to reduce with dithiothreitol. These experiments proved that Kcnq4 is present in DRG but not in keratinocytes and is indeed a direct molecular target of H_2_O_2_-mediated redox changes.

## Discussion

To our knowledge, the present work is the first to demonstrate the physiological sensory function of NADPH oxidase-derived ROS in the mammalian skin. We analyzed the expression and activity of the members of the NOX/DUOX family of NADPH oxidases in the mouse skin and primary keratinocytes as well as in human immortalized keratinocytes and showed that the predominant member of this family in both mouse and human keratinocytes was DUOX1. We generated a specific, polyclonal antibody against DUOX1 so that we could also confirm its expression at the protein level. We also devised a technique to visualize its hydrogen peroxide-producing activity in epithelial monolayers.

DUOX1 is a transmembrane NADPH oxidase enzyme possessing intracellular Ca^++^-binding EF-hands and its activity can be induced through intracellular Ca^++^ signals. It also carries an extracellular peroxidase homology domain which is inactive in mammalian species. Despite a large number of publications and the recently published detailed structural information about DUOX1 (Wu et al., 2021), relatively little is known about the physiological functions and biochemical mechanisms of this enzyme. Previously it has been described in various functions, like the airway epithelial wound repair and epithelial responses to allergic and microbial triggers (Boots et al., 2009; Hristova et al., 2016; Sarr et al., 2021; Wesley et al., 2007), the pressure responses of the urinary bladder (Donkó et al., 2010), the activity of primary B-cells (Sugamata et al., 2019) or the growth factor elicited redox signaling of epithelial cells (Sirokmány et al., 2016). Interestingly, epigenetic downregulation of DUOX1 was reported in different epithelial cancers (Ling et al., 2014; Little et al., 2016; Luxen et al., 2008). It has also been shown that DUOX (Ce-DUOX1/BLI-3) plays an important role in the development and maintenance of the *C. elegans* cuticle (Edens et al., 2001; Kamath et al., 2003). Importantly, however, we have not found any structural or ultrastructural differences in the skin of Duox1 deficient mice.

As DUOX1 activity can be triggered by various stimuli in keratinocytes it seemed to be a plausible hypothesis that environmental inputs can affect the sensory function of the skin by altering the DUOX1 oxidase activity. This possibility seemed particularly interesting because keratinocytes are increasingly being recognized as active participants in sensory functions.

It has been shown previously that some molecules, like the Transient Receptor Potential Vanilloid 3 (TRPV3) can play a role in both sensory and barrier functions of the skin (Cheng et al., 2010; Huang et al., 2011b; Moqrich et al., 2005). As the proper development of the barrier function of the skin can also severely affect its sensitivity towards environmental challenges, it was requisite to thoroughly investigate possible morphological changes in DUOX1 knockout skin. Our detailed histological analysis indicates that there are no structural or developmental defects in DUOX1- deficient mammalian skin.

We found an enhanced response of DUOX1-knockout animals towards thermal nociceptive treatments following allyl-isothiocyanate pretreatment. DUOX1-knockout animals also showed increased nocifensive responses after intraplantar formalin injection. In this regard, it is also a key result of our gene expression analysis that DUOX1 is not expressed in sensory neuronal cells. Importantly we found no difference in mechanonociceptive thresholds. The selective difference in the nocifensive behavior towards thermal stimuli also indicated an altered sensitivity of peripheral nerve endings of the skin. In agreement with these findings, mechanical hyperalgesia is reported to be under different central control than thermal hyperalgesia (Otsubo et al., 2012). As DUOX1 is expressed specifically in epidermal keratinocytes but not in neuronal cells our finding provides a possible therapeutic target to manage peripheral pain conditions without a direct impact on central nervous functions.

We propose that DUOX1 acts as a quickly and reversibly inducible signaling center, the H_2_O_2_ production of which can influence direct molecular targets in paracrine and/or autocrine fashion. The autocrine way of H_2_O_2_ action could be the regulation of the release of signaling molecules from keratinocytes. There is a large number of mediators that had been reported to be released from keratinocytes. Adenosine triphosphate, prostaglandins, leukotrienes, TNF-alpha, interleukins, and endogenous opioids have all been reported to be released from keratinocytes (Andoh et al., 2001; Ansel et al., 1990; Bigliardi et al., 2009; Burrell et al., 2005). Out of these, we tested PGE_2_ and ATP release, both well known for eliciting nociceptive responses, but we found no differences in DUOX1-deficient keratinocytes. Obviously, we cannot exclude the possibility that the secretion of some other mediators might be affected by DUOX1 activity.

In comparison with the above-described signaling molecules, H_2_O_2_ has got several favorable properties. It can be produced very quickly, and its secretion cannot be depleted. It has a very small molecular size which enables quick diffusion in the extracellular space. Finally, there are several enzymatic and non-enzymatic ways to quickly inactivate or metabolize it which contributes to tight regulation of its biological activity. To delineate DUOX1-derived ROS as a paracrine mediator of nociceptive signaling we aimed at identifying possible molecular targets of hydrogen peroxide. We browsed literature data of cell surface proteins expressed on sensory nerve fibers that were reported to display redox sensitivity. We selected TRPA1 and KCNQ4 for further analysis.

In accordance with previous data, H_2_O_2_ could elicit calcium signals in TRPA1-expressing cells. We found that pretreatment of TRPA1 expressing cells with H_2_O_2_ resulted in decreased responsiveness to TRPA1 agonists. This suggests a desensitizing action of H_2_O_2_ on sensory nerves, damping the general reactivity to nociceptive stimuli. Another way to frame this effect of H_2_O_2_ is that it actually decreases the contrast between the non-stimulated and AITC-stimulated conditions and thereby lowers the S/S_0_ ratio in the Weber-Fechner equation. This leads to the increase of the just noticeable difference, the smallest change in stimuli that can be perceived.

We also show biochemical evidence of H_2_O_2_-mediated KCNQ4 oxidation with as low as 20 μM H_2_O_2_. Currently, we do not know if KCNQ4 and TRPA1 are present on the same or rather on some different DRG cell populations. The development of specific antibodies suitable for immunohistochemistry will improve our understanding in this regard. However, the slower closure of KCNQ4 channels following exposure to H_2_O_2_ might also contribute to a slower or reduced peripheral activation of sensory nerve fibers.

It is also very interesting to note that our previous study has demonstrated that DUOX1 plays a regulatory role in the mechanosensing function of urothelial epithelia. The lack of DUOX1 activity in knockout animals resulted in an enhanced frequency of voiding contractions of the bladder (Donkó et al., 2010). Although this is a different physiological setting compared to cutaneous nociception, it is intriguing that also in this mechanosensory function DUOX1 had a dampening, alleviating effect. In summary, our results shed light on a hitherto unrecognized physiological sensory function of DUOX1 in the skin. We pinpointed potential molecular targets of the DUOX1-derived hydrogen peroxide. We believe that H_2_O_2_-mediated effects include - but are probably not limited to - these molecules. Our data suggest a subtle, physiological regulatory role for ROS in sensory functions. This is rather in contrast to the previous conception of ROS as a pain enhancer factor (Salvemini et al., 2011; Wang et al., 2004). The main difference in the interpretation of the role of ROS lies probably in the source and regulation of ROS release. Robust ROS production by tissue invading leukocytes in the setting of a tissue injury or severe inflammation is very different from the continuous, small-scale, physiological ROS production of keratinocytes in the intact epidermis.

Our results about ROS released from keratinocytes might open a new path to a more detailed understanding of nociceptive and analgesic signaling pathways in the skin. Further analysis of molecules affected by keratinocyte-derived H_2_O_2_ can provide actionable molecular targets.

This can contribute to local treatments of pain states without the side effects of systemic therapy.

## Methods

### Cell culture

HEK293 (ATCC, Manassas, VA, USA, CRL-1573), HEK293T (ATCC, Manassas, VA, USA, CRL-3216), HEK293A (Invitrogen R705-07) and HaCaT cells (ATCC, Manassas, VA, USA) were cultured in Dulbecco’s Modified Eagle’s Medium (DMEM; Lonza Group Ltd., Basel, Switzerland) supplemented with 100 U/ml penicillin, 100 U/ml streptomycin (Lonza Group Ltd., Basel, Switzerland) and 10% fetal bovine serum (FBS; Lonza Group Ltd., Basel, Switzerland). Cells were grown in a humidified incubator with 5% CO_2_ in air, at 37 °C.

Primary mouse keratinocytes were prepared from the back skin of wild-type and Duox1 deficient mice as described previously (Romero et al., 1999).

### Animals

*DUOX1* knockout mice described earlier by Donkó *et al* (Donkó et al., 2010) were purchased from Lexicon Pharmaceuticals, Inc. (The Woodlands, TX, USA). C57BL/6N wild-type mice were obtained from commercial sources. Experimental animals were maintained on a standard diet and given water *ad libitum* in either specific pathogen-free or conventional animal facilities of Basic Medical Science Centre, Semmelweis University.

Animals for nociception assays were kept in the animal house of Szentágothai Research Centre, University of Pécs, in individually-ventilated cages on wood shavings bedding, at 22±2 °C temperature and a standard 12-12 h light-dark cycle (lights on between 7:00-19:00). They were provided with standard rodent chow and water *ad libitum*. All experiments were designed and performed according to the 243/1998. Hungarian government regulation on animal experiments. Experiments were approved by the Ethics Committee on Animal Research of the University of Pécs (license number: BA02/2000-10/2011).

10-12 week old male mice of each genotype were used up to the necessary sample size. Sample size was determined based on our previous experiments using the same models (Tékus et al., 2016) considering also the 3R (replace, reduce, refine) rule for the animal ethical principles. The animals were randomized in the different experimental groups, the researchers were blinded to the experimental design, the treatment the animals received, and the genotype. Due to short-term anaesthesia and short observation times, the general health status of animals was not affected. The analysis included the results of all tested animals, there were no exclusions.

### Mustard oil-induced thermal hyperalgesia on the tail

The noxious heat threshold of the tail of mice was measured with an increasing-temperature water bath (Experimetria Ltd., Budapest, Hungary) as previously described (Tékus et al., 2016). Mice were placed into plastic restrainers which were hanged on a rack above the water bath so that the tails could be immersed into the water. The following parameters were set on the device: a starting temperature of 30 °C, a heating rate of 24 °C/min and a cut-off temperature of 53 °C. The heating was immediately stopped when the animals showed nocifensive behavior and the tails were removed from the water. The corresponding water temperature was recorded as the noxious heat threshold. After 3 control treshold measurements, thermal hyperalgesia was induced by immersing the tails into 5% mustard oil dissolved in 30% DMSO in water for 30 seconds. Noxious heat threshold measurements were repeated at 10 min intervals for 60 minutes after treatment. Where indicated, 3mg/kg intraperitoneal naloxone pretreatment had been applied.

### Heat injury-induced thermal and mechanical hyperalgesia on the paw

The noxious heat threshold of the paws of freely-moving mice was measured with an increasing-temperature hot plate (IITC Life Sciences, Woodland Hills, CA, USA), while the mechanonociceptive threshold was determined by a dynamic plantar aesthesiometer (DPA; Ugo Basile, Comerio, Italy). The starting temperature on the hot plate was set to 30 °C and a heating rate of 12 °C/min and cut-off temperature of 53 °C were used. For the mechanical stimulation, the DPA device uses a blunt needle pushed at the plantar surface of the paw at an increasing force, which was set to 2 g/sec up to a cut-off value of 10 g. For both methods, stimulation was immediately stopped when the animal showed nocifensive behavior and the corresponding temperature or force was recorded as the threshold. After 3 control threshold measurements, heat injury (i.e. a first-degree burn model) was induced under ether anaesthesia by immersing one of the hind paws into 51 °C water for 15 seconds. Noxious thermal or mechanical thresholds were measured for 60 minutes at 10 min intervals afterwards (Bölcskei et al., 2005).

### Plantar incision-induced mechanical hyperalgesia on the paw

Mechanical thresholds were measured by the DPA, as described above. A postoperative pain model was induced by making a 5-mm incision on the paw, under ketamine-xylazine anaesthesia (100-5 mg/kg i.p.). The incision was closed with 4/0 silk sutures and animals were allowed to recover for 24 h. Thresholds were measured afterwards once per day for 3 days after surgery (Pogatzki and Raja, 2003).

### Formalin test

For the evaluation of formalin-induced nociception, mice were injected intraplantar with 2.5% formalin (20 μl, i.pl.) and then placed into transparent observation chambers. A mirror was placed behind the chamber so that the observer could see the animals from behind. Spontaneous nocifensive behavior was assessed between 0-5 min and 20-45 min after injection, based on the characteristic 2-phase response induced by formalin. The duration of paw licking was measured by a stopwatch, while paw flinching responses were counted. A Composite Pain Score (CPS) was also calculated by the following formula: CPS=(2x paw licking time + 1x paw flinchings)/observation time. The swelling induced by formalin was determined 3 h after injection by measuring the thickness of the paw with a digital caliper (Mitutoyo, Kawasaki, Japan). The swelling was expressed as a % increase compared to the contralateral paw.

### Histology and immunostaining

Tissues were fixed in formal saline and embedded in paraffin blocks. For routine histology and immunostaining, 5 μm thick sections were cut. These were either stained with haematoxylin and eosin or went through heat induced epitope retrieval (HIER) in citrate buffer (10 mM sodium citrate, pH 6.0) before overnight incubation with specific first antibodies. To be able to detect PGP9.5-positive nerve fibers in the epidermis, 100 μm thick paraffin sections were cut. These were also processed through HIER and the floating thick skin sections were incubated overnight with rabbit polyclonal anti-PGP9.5 antibodies. Multiple Z-stack images acquired on a Leica SP5 laser scanning microscope, using a 20x HCX PL APO CS dry objective were merged to obtain the final images. For Tuj-1 wholemount labeling tail skin was slit lengthways with a scalpel, peeled off and cut into 5×5 mm pieces, and incubated in 5 mM EDTA in PBS at 37 °C for four hours. The intact sheet of epidermis was gently peeled away from the dermis and the epidermal tissue was fixed in 4% formal saline for 2 hours at room temperature (Braun et al., 2003). Fixed epidermal sheets were blocked, permeabilized and incubated overnight with Tuj-1 antibody. After extensive washing steps (6x 30 min), the secondary fluorescent antibody was added for four hours, and after further washing steps, the epidermal tissue pieces were mounted on glass slides.

Primary antibodies: keratin-10, keratin-14, loricrin and gamma-catenin antibodies were from Covance Research Products, Inc. (now BioLegend). Connexin-43 antibody (#3512) was from Cell Signaling Technology, Danvers, Massachusetts, USA, Tuj-1 antibody (sc-58888) was from Santa Cruz Biotechnology Inc, Dallas, Texas, USA. PGP9.5 antibody (ab8189) was from Abcam, Cambridge, UK. Alexa488 or Alexa568 coupled secondary antibodies were from ThermoFisher Scientific.

### Electron microscopy

Sample preparation and imaging were carried out as described previously (Kollo et al., 2006; Lörincz et al., 2002). Briefly, adult mice were anesthetized with ketamine (0.4 ml/100g, 0.2 ml/mouse i.p.), then were transcardially perfused with 0.9 % saline for 1 min followed by the fixative: 4% paraformaldehyde, 15% picric acid, 0.05% glutaraldehyde made up in 0.1 M phosphate buffer for 30 min. Sections for electron microscopy were postfixed with 0.5-1% OsO_4_, contrasted in 1% uranyl acetate, dehydrated in graded alcohol series, and embedded into epoxy resin. Sections were examined with a transmission electron microscope (TEM, JEM1011, Jeol).

### Quantitative PCR

Mouse tissue RNA was isolated from dorsal root ganglions (DRG), hindpaw and tail skin using RNeasy Mini Kit (Qiagen, Hilden, Germany). Human keratinocyte RNA was purified using NucleoSpin RNA (Macherey-Nagel, Düren, Germany). Before the RNA preparation, tissue sample were collected into RNAlater reagent (Thermo Fisher Scientific, Waltham, MA, USA) at room temperature. The concentration of the RNA samples was tested by ultraviolet absorption at 260/280 nm in Nano Drop One system (Thermo Fisher Scientific, Waltham, MA, USA). cDNA was synthesized from 2 μg of total RNA using High-Capacity cDNA Reverse Transcription Kit (Fermentas) according to the manufacturer’s recommendations. For qPCR reaction 0.5 μl of cDNA was used in a 10 μl reaction solution using Taqman Gene Expression Assays (Thermo Fisher Scientific, Waltham, MA, USA) and LightCycler 480 Probes Master (Roche Life Science) in a LightCycler LC480 plate reader (Roche Life Science). For each cDNA sample, the expression of the target was divided by the expression of the endogenous control, which was *eukaryotic elongation factor 2* (Eef-2) or GAPDH. The crossing point was determined by the second derivative method.

The following Taqman Gene Expression assays were used:

#### Mouse

Nox1: Mm00549170_m1, Nox2: Mm01287743_m1, Nox4: Mm00479246_m1, DUOX1: Mm01328698_m1, Duox2: Mm01326247_m1, DuoxA1: Mm01269313_g1, DuoxA2: Mm00470560_m1, Eef-2: Mm01171434_g1, Kcnq4: Mm01185500_m1, TrpA1: Mm01227437_m1, TrpV1: Mm01246302_m1, TrpV3: Mm00455003_m1, TrpV4: Mm00455003_m1, Dpp6: Mm00456620_m1, DPP10: Mm01284949_m1, Kcnd3 Mm01302126_m1, Kcnd1: Mm00492793_g1, GAPDH: Mm99999915_g1.

#### Human

Nox1: Hs00246589_m1, Nox2: Hs00166163_m1, Nox3: Hs01098883_m1, Nox4: Hs00418356_m1, Nox5: Hs00225846_m1, DUOX1: Hs00213694_m1, Duox2: Hs00204187_m1, DuoxA1: Hs00328806_m1, DuoxA2: Hs01595311_g1 p22phox: Hs03044361_m1, NoxO1: Hs00376045_g1, NoxA1: Hs00736699_m1, GAPDH: Hs99999905_m1.

### Amplex Red assay

Confluent cells on 24-well plates (SPL Life Sciences Co., Korea) were washed with H-medium (145 mM NaCl, 5 mM KCl, 1 mM MgCl_2_, 0.8 mM CaCl_2_, 10 mM HEPES, and 5 mM glucose, pH 7.4) and background fluorescence was also measured in 0.3 ml/well H-medium. For the assay, an H-medium-based reaction solution was used containing horse radish peroxidase (Sigma-Aldrich, Burlington, MA, USA) and Amplex Red (Synchem, Germany) in a final concentration of 0.2 U/ml and 50 μM respectively. Cells were stimulated with 1 μM thapsigargin (Sigma-Aldrich, Burlington, MA, USA) or 10 μM ATPγS (Sigma-Aldrich, Burlington, MA, USA), or 2 nM GSK 1016790A (Sigma-Aldrich, Burlington, MA, USA). Stimuli were also added to the Amplex Red containing reaction solution immediately before pipetting it onto the cells. The measurement of fluorescence started promptly after the addition of the reaction solution and the cells were kept at 37 °C throughout the measurement. Fluorescence of the end-product resorufin was measured at 590 nm with POLARstar Optima multidetection microplate reader (BMG Labtech, Ortenberg, Germany). Background fluorescence was subtracted from the fluorescence values of each well. Each experimental condition was run in 3 parallels on the 24-well plate.

### Measurement of Prostaglandin E_2_ release

SiRNA-treated HaCaT cells were stimulated with ATPγS and cell culture supernatants were harvested after the indicated incubation times. PGE_2_ was measured using the Enzo Life Sciences PGE_2_ ELISA kit, according to the manufacturer’s instructions. Concentrations were calculated based on a calibration curve.

### Measurement of ATP secretion

HEK293A cells expressing the GRAB_ATP_ sensor were kindly provided by Balázs Enyedi. GRAB_ATP_1.0 expression plasmid was created by Gibson cloning based on the sequence provided in the article of Wu et al. (Wu et al., 2022). GRAB_ATP_1.0 was N-terminally fused through a P2A peptide with a plasma membrane localized far-red fluorescent protein, mKate2. Membrane localization of mKate2 was achieved by fusing it to the N-terminal targeting sequence of the protein Lck (MGCVCSSNPENNNN).

35000 cells/well (30% HEK293A-70% HaCaT or 100% HEK293A) were plated on Ibidi 8-well μ-slides (Ibidi GmbH, Gräfelfing, Germany) pretreated with poly-L-lysine (Sigma-Aldrich, Burlington, MA, USA). Next day before measurements, growth medium was replaced with imaging medium (EC1, containing 3.1 mM KCl, 133.2 mM NaCl, 0.5. mM KH2PO4, 0.5 mM MgSO4, 5 mM Na-HEPES, 2 mM NaHCO3, 1.2 mM CaCl2 and 2.5 mM Glucose) Experiments were performed at room temperature on a NikonTi2 inverted microscope equipped with an Apo LWD 40x WI λS DIC N2 water immersion objective, a Yokogawa CSU-W1 Spinning Disk unit, a Photometrics Prime BSI camera and 488 nm and 561 nm diode laser lines. After recording the cells for 2 minutes, a custom-made perfusion system was opened and fresh EC1 medium was delivered by perfusion to wash the previously produced ATP away. As treatment 2.5 nM GSK 1016790A or 500 μM H_2_O_2_ were applied. At the end of the measurement 1 μM ATP was added as a positive control.

### Analysis and quantification of ATP secretion

GRAB_ATP_ expressing cells were automatically segmented in the red channel (mKate2) using the software Cellpose (Stringer et al., 2021). The mean intensity of the green and red channel (cpEGFP and mKate2) was recorded for each cell at each time point using the generated masks. Taking advantage of the P2A peptide linker between the mKate2 and GRAB_ATP_ resulting in a 1:1 expression ratio, we used the green/red ratio as a normalized measure of the GRAB_ATP_ signal. To express all intensities between 0 and 1, we applied the following further normalization: (F-F_0_)/(F_max_-F_0_), where F is the intensity at a given time point, F_0_ is the average intensity value of the baseline measured during the 5 minutes prior to the stimulation and F_max_ is the maximum intensity value after ATP stimulation. Finally, we applied a rolling average of 3 on the normalized data. The final output data was then denoted ΔF/F_0_. For all experiments, we excluded the 2 minute pre-prefusion baseline and the first 3 minutes of the washing step from the analysis.

Prior to analysis, the background intensity was removed automatically using the SMO software (https://doi.org/10.1101/2021.11.09.467975) and images were registered using the pystackreg software (https://doi.org/10.1109/83.650848). All data were analyzed and plotted using the Python libraries

Pandas https://doi.org/10.25080/MAJORA-92BF1922-00A

Numpy https://doi.org/10.1038/s41586-020-2649-2

Seaborn https://doi.org/10.21105/joss.03021

Statistical analysis was done using the t-test of the ‘stats’ module from the Python library Scipy.

### Calcium imaging

40000 cells/well were plated on 96-well plates (SPL Life Sciences Co., Korea) pretreated with poly-L-lysine (Sigma-Aldrich, Burlington, MA, USA). HEK293 cells were transfected with human TRPA1 expressing plasmid using Lipofectamine 2000 (Invitrogen) according to the manufacturer’s instructions. The plasmid was kindly provided by Zoltán Sándor, Department of Pharmacology and Pharmacotherapy, University of Pécs, Medical School (Pozsgai et al., 2017). Next day cells were washed with H-medium and loaded with Fura2-AM (2 μM dissolved in H-medium, Molecular Probes, USA) for 30 min at 37 °C. Cells were washed again and the baseline fluorescence was measured using Clariostar (BMG Labtech, Ortenberg, Germany). Excitation wavelengths were 335 and 380 nm, emission wavelength was 510 nm. The final concentrations of different stimuli were as follows: 500 or 100 μM H_2_O_2_ (Sigma-Aldrich, Burlington, MA, USA) and 10 μM AITC (Sigma-Aldrich, Burlington, MA, USA). Each experimental condition was run in 3 parallels.

### OIKBiotinylation of reduced thiols

HEK293T cells were transfected on poly-L-lysine (Sigma-Aldrich, Burlington, MA, USA) coated 6-well plates (SPL Life Sciences Co., Korea) with Kcnq4 Mouse Tagged ORF Clone (OriGene, Rockville, MD, USA) using Lipofectamine LTX and Plus Reagents (Invitrogen) following the manufacturer’s instructions. Next day the transfected cells were washed with H-medium and treated with 0, 20, 100, or 500 μM H_2_O_2_ in 2 ml H-medium for 3 minutes at 37 °C. After the treatment cells were lysed and collected in ice-cold lysis buffer (50 mM TRIS, 140 mM NaCl, 1% Triton X-100, 0.1% SDS, 1mM PMSF, and cOmplete Mini Protease Inhibitor Cocktail (Merck, Darmstadt, Germany), Ph 8) with 250 μM biotin polyethyleneoxide iodoacetamide (BIAM, Thermo Fisher Scientific, Waltham, MA, USA). Samples were centrifuged at 15,000 RPM for 10 minutes at 4°C and the supernatants were incubated for 25 min at 4 °C. The protein concentration of the samples was adjusted to equal amounts using Pierce BCA Assay Kit (Thermo Fisher Scientific, Waltham, MA, USA) for subsequent immunoprecipitation. 50-100 μg of protein were used for the pull down of FLAG-tagged Kcnq4 proteins with monoclonal anti-FLAG M2 antibody produced in mouse (mouse monoclonal Anti-FLAG M2 antibody, F3165) and Protein G Sepharose beads (Abcam, UK) at 4 °C overnight. For Western blot analysis, beads were boiled in Laemmli sample buffer.

### Biotinylation of reversibly oxidized thiols

The experiment started like the biothinylation of reduced thiols, described above. After H_2_O_2_ treatment, biotin labeling of reversibly oxidized thiols was conducted by a modified protocol based on Oliver Löwe et al.(Löwe et al., 2019). Briefly, free thiols were alkylated with 100 mM N-ethylmaleimide (NEM, Sigma-Aldrich, Burlington, MA, USA) in H-medium for 5 minutes at room temperature. Subsequently, cells were lysed and collected from the plate in ice-cold BASE-buffer (1% IGEPAL-CA630, 150 mM NaCl, 50 mM TRIS, 1 mM EDTA, pH 8) supplemented with 100 mM NEM and cOmplete Mini Protease Inhibitor Cocktail (Merck, Darmstadt, Germany) on ice. Samples were centrifuged at 15,000 RPM for 10 minutes at 4 °C. Excess of NEM was removed from the supernatants using 0.5 ml Zeba Spin Desalting Columns (Thermo Fisher Scientific, Waltham, MA, USA) according to the manufacturer’s recommendations. Reversibly oxidized thiols were reduced with 2.3 mM dithiothreitol (DTT, Avantor, Radnor, PA, USA) for 30 minutes on ice. Upon removal of excess DTT with desalting columns, reduced thiols were alkylated with biotin polyethyleneoxide iodoacetamide (Thermo Fisher Scientific, Waltham, MA, USA) for 2 hours on ice in ultrasound bath. After removal of unbound BIAM with desalting columns, the steps of the biothinylation of reduced thiols as described above were followed. Kcnq4 was immunoprecipitated and signals were detected by western blot.

### Biotinyl tyramide assay

HaCaT and HaCaT DUOX1 knockout confluent cells on coverslips were washed with H-medium and treated with 1 μM thapsigargin in the presence or absence of 0.2 U/ml horse radish peroxidase and 27,5 μM biotinyl tyramide (Sigma Aldrich, Burlington, MA, USA) for 5 minutes at 37 °C. After treatment, cells were washed 3 times with PBS on ice and bound biotinyl tyramide was visualized with fluorescent streptavidin (Vector Laboratories, Inc., Burlingame, CA, USA) at 1:1000 in PBS for 30 minutes at 4 °C. Cells were fixed with ice-cold 4% paraformaldehyde solution and cell nuclei were stained with To-Pro-3 (Invitrogen). Samples were mounted with Mowiol (Sigma Aldrich, Burlington, MA, USA) and analyzed with an LSM710 confocal laser-scanning microscope using a 63x oil objective (Carl Zeiss).

### Non-commercial DUOX1 antibodies

411-amino-acids-long sequence of recombinant human DUOX1 (amino acids 622-1032) was produced and purified from BL21 Competent Cells and injected intracutaneously with Freund’s adjuvant into New Zealand white rabbit.. For polyclonal antibodies rabbits were sacrificed, and antibodies were affinity purified from the sera using Affigel 10 beads (BioRad Laboratories, Hercules, CA, USA) loaded with the antigens according to the manufacturer’s instructions.

### DUOX1 CRISPR in HaCaT

HaCaT cells were genetically mutated for DUOX1, using a pSpCas9(BB)−2A-GFP (PX458, Addgene) vector, following Target Sequence Cloning Protocol by ZhangLab. The vector contained the 5′-gagctgtctcggctgcggacagg-3′ guide sequence. The cells were transfected with Lipofectamine LTX and Plus Reagent (Invitrogen) and GFP-positive cells were sorted onto 96 well plates. Cell clones were screened by PCR of genomic DNA using 5′-gtgcagtgaggatatcccaaccc-3′ sense and 5′-ctggctcctgaccaatgctgg-3′ antisense oligos. PCR products were analyzed by Surveyor mismatch analysis, then sent for sequencing. It was confirmed by Western blot analysis and Amplex Red assay that the selected cell line does not express DUOX1.

### Western blot experiments

Laemmli sample buffer was added to the cell lysate samples and these were run on 8% or 10% SDS polyacrylamide gels and blotted onto nitrocellulose membranes. Membranes were blocked in phosphate buffered saline containing 0,1% Tween-20 and 5% dry milk. The first antibodies were diluted in blocking buffer and used either for 1 h at room temperature or overnight at 4 °C. After several washing steps in PBS-Tween-20 membranes were incubated with HRP-linked secondary antibodies (Amersham Pharmaceuticals, Amersham, UK) diluted in blocking buffer. After further PBS-Tween-20 washing steps antibody binding was detected using enhanced chemiluminescence and Fuji Super RX medical X-ray films. Importantly samples were never boiled when processed for western blotting with the DUOX1 antibody.

The redox state of Kcnq4 was also measured by immunoblotting. Detection of proteins was performed using HRP-Conjugated Streptavidin (Thermo Fisher Scientific, Waltham, MA, USA) in phosphate buffered saline containing 0,1% Tween-20 and 5% bovine serum albumin.

### Statistical analysis

Statistical analyses were performed using Graph Pad Prism 7.0 and Origin Pro 8 software programs. Specific statistical tests are presented in the figure legend for each experiment. *P-*values below 0.05 were considered statistically significant.

## Acknowledgements

We are grateful to Beáta Molnár, Barbara Bodor-Kis, Dóra Ömböli and Dóra Rónaszéki for technical assistance. We also appreciate the expert help of Zoltán Nusser in electron microscopy. We are thankful for the work of the animal facility technicians Klára Papp and Àdám Marinkás. This study was supported by grants from the National Research, Development and Innovation Office (PD138404, K133002, NVKP_16-1-2016-0039). The work was also financed by the Thematic Excellence Program 2021 Health Subprogram (MOLORKIV) of the Ministry for Innovation and Technology in Hungary and by grants VEKOP-2.3.2-16-2016-00002 and EFOP-3.6.3-VEKOP-16-2017-00009. This project was also supported by the National Research, Development and Innovation Office (PharmaLab, RRF-2.3.1-21-2022-00015), Eötvos Loránd Research Network, TKP2021-EGA-16 has been implemented with the support provided from the National Research, Development and Innovation Fund of Hungary, and OTKA K-138046.

A.P. was supported by a predoctoral grant EFOP-3.6.3-VEKOP-16-2017-00009. B. Enyedi was supported by a “Lendület” grant from the Hungarian Academy of Sciences (LP2018-13/2018) and funding form EU’s Horizon 2020 research and innovation program (grant agreement No. 739593).

---

**Figure 3 - figure supplement 1.**
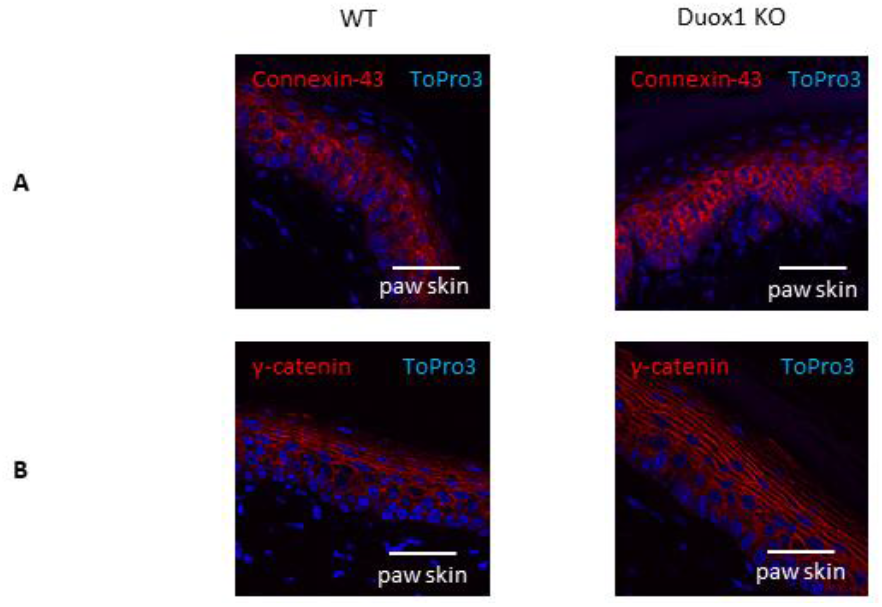
Histological analysis of wild-type and DUOX1-deficient mouse skin. (*A* and *B*) Wild-type or DUOX1 KO mouse paw skin sections were labeled with connexin-43 (*A*) and γ-catenin antibodies (*B*). Scale bars: 25 μm.

**Figure 3 - figure supplement 2.**
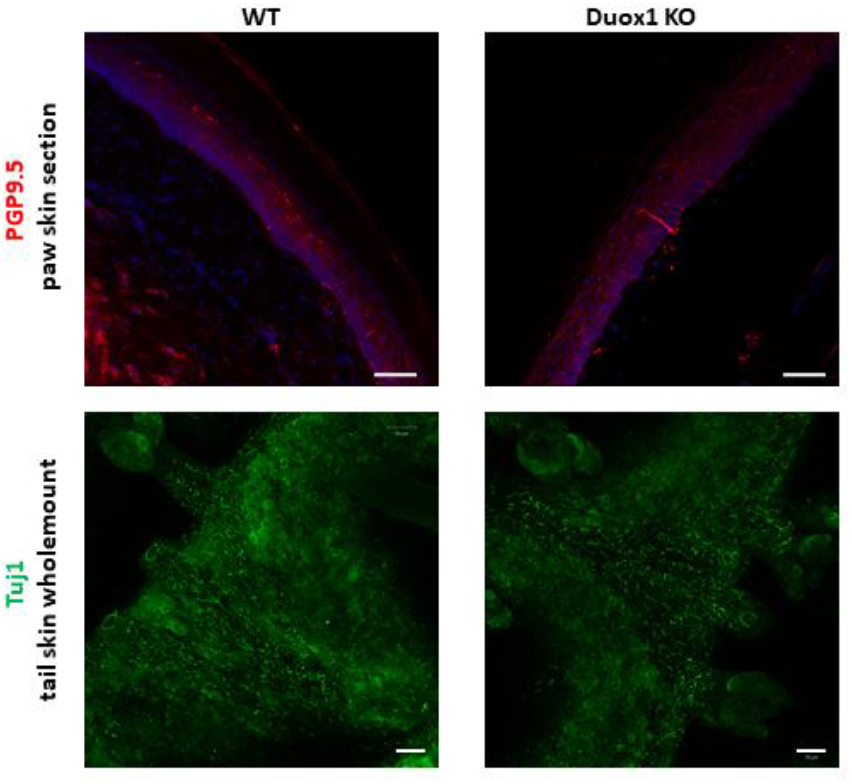
Immunolabeling of epidermal nerve fibers of mouse paw and tail skin. (*A*) 100 μm thick paraffin sections were labeled with anti-PGP9.5 antibody and DAPI nuclear stain. Scale bars: 25 μm. (*B*) Tuj1 was stained on tail skin wholemount tissue samples. Scale bars: 50 μm.

**Figure 4 – figure supplement 1.**
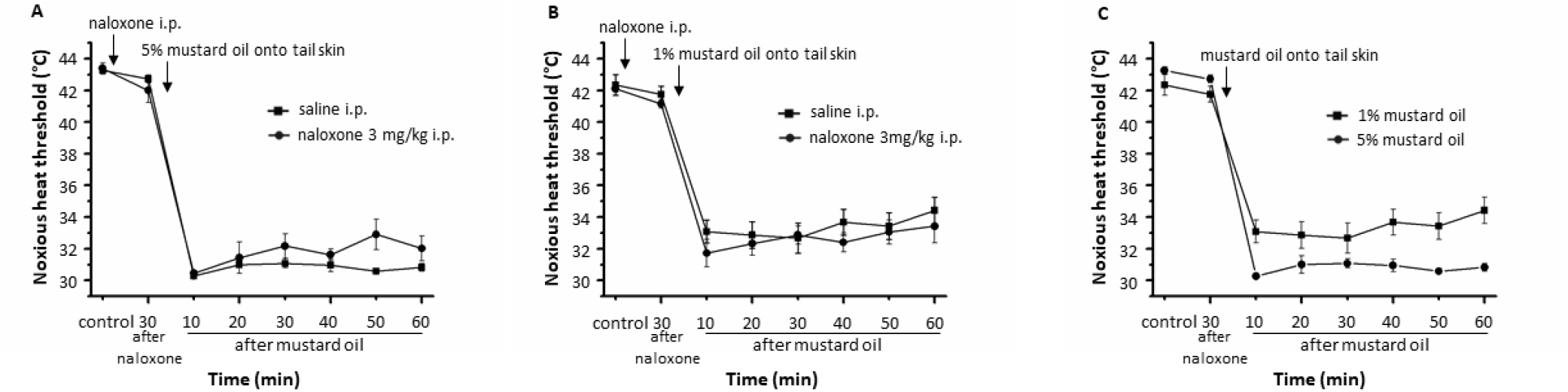
Measurement of thermal nociceptive threshold after naloxone injection. (*A, B* and *C*) Noxious heat threshold of the tail measurement was carried out after 3 mg/kg intraperitoneal naloxone injection. Thermal hyperalgesia was induced by immersing the tails into 5% (*A*) or 1% (*B*) mustard oil. Plots show mean ± SEM of n=6 animals/group.

**Figure 5 - figure supplement 1.**
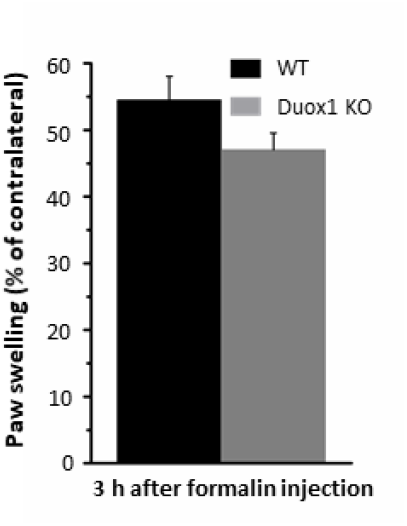
Formalin-induced nociception in wild-type and Duox1 knockout mouse. The swelling induced by formalin was determined 3 h after injection by measuring the thickness of the paw. The swelling was expressed as a % increase compared to the contralateral paw. Data are mean ± SEM of n=5-5 animals/group.

**Figure 5 - figure supplement 2.**
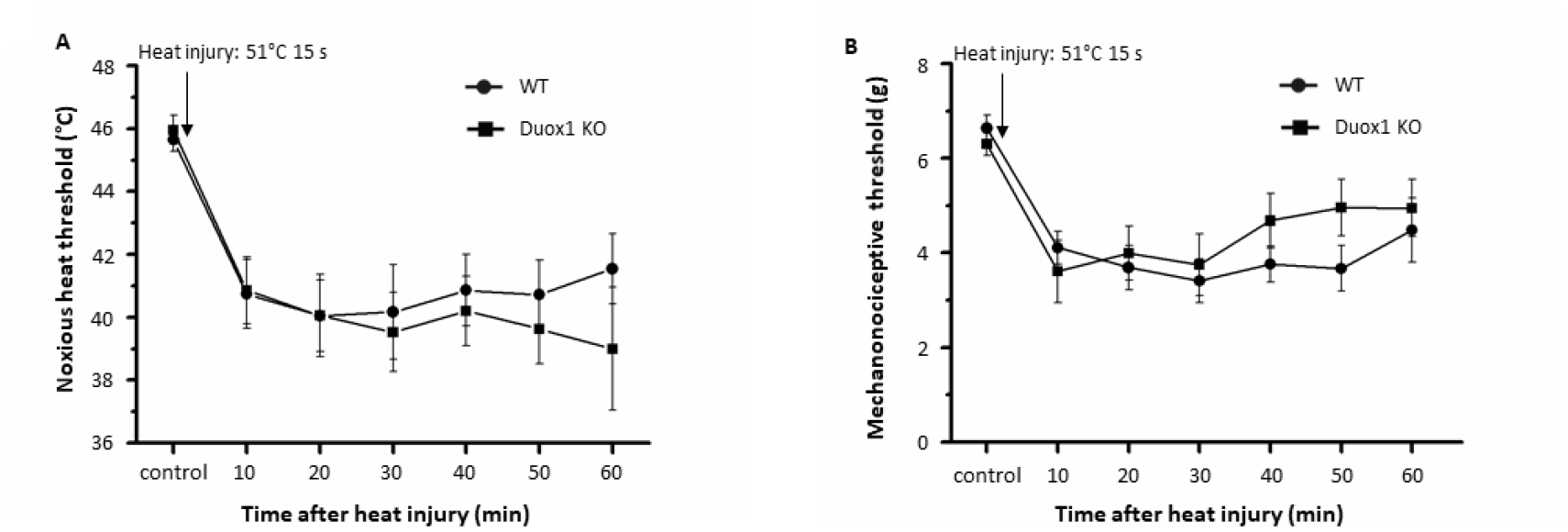
Heat injury-induced thermal and mechanical hyperalgesia on the paw. Heat injury was induced under ether anaesthesia by immersing one of the hind paws into 51°C water for 15 seconds. (*A*) Noxious heat threshold of the paws of freely-moving mice was measured with an increasing-temperature hot plate. (*B*) Mechanonociceptive threshold was determined by a dynamic plantar aesthesiometer (DPA). Noxious thermal or mechanical thresholds were measured for 60 minutes every ten minutes. Plots show mean ± SEM of n=10 animals/group.

**Figure 5 - figure supplement 3.**
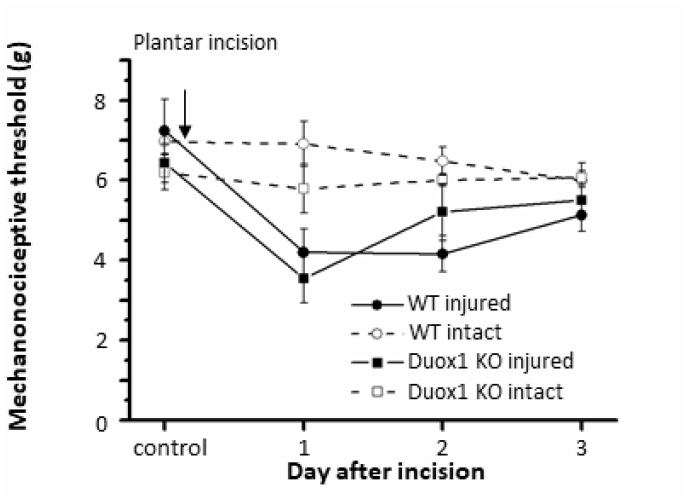
Plantar incision-induced mechanical hyperalgesia on the paw. A postoperative pain model was induced by making an incision on the paw, under ketamine-xylazine anaesthesia. Noxious mechanical thresholds were measured afterwards once per day for 3 days after surgery by DPA. Data are mean ± SEM of n=5-5 animals/group.

**Figure 6 – figure supplement 1.**
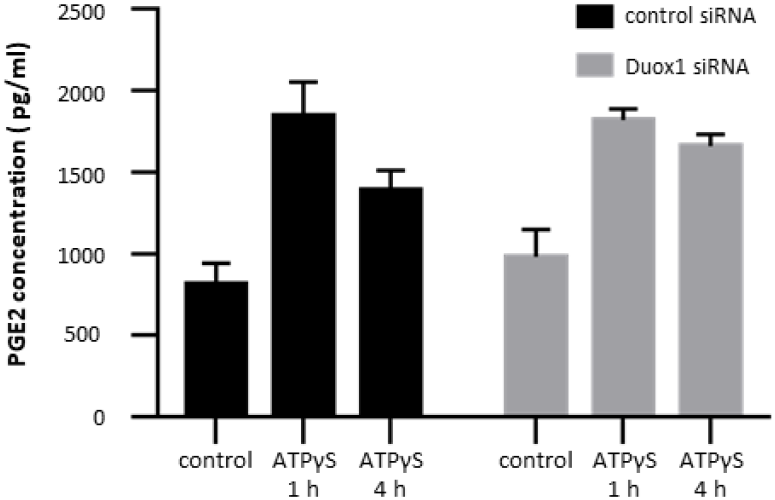
ATPγS-induced PGE2-release of scrambled or DUOX1-specific siRNA treated HaCaT cells. Representative plot shows mean ± SD of triplicate. The experiment was repeated twice, with similar results.

**Figure 6 – figure supplement 2.**
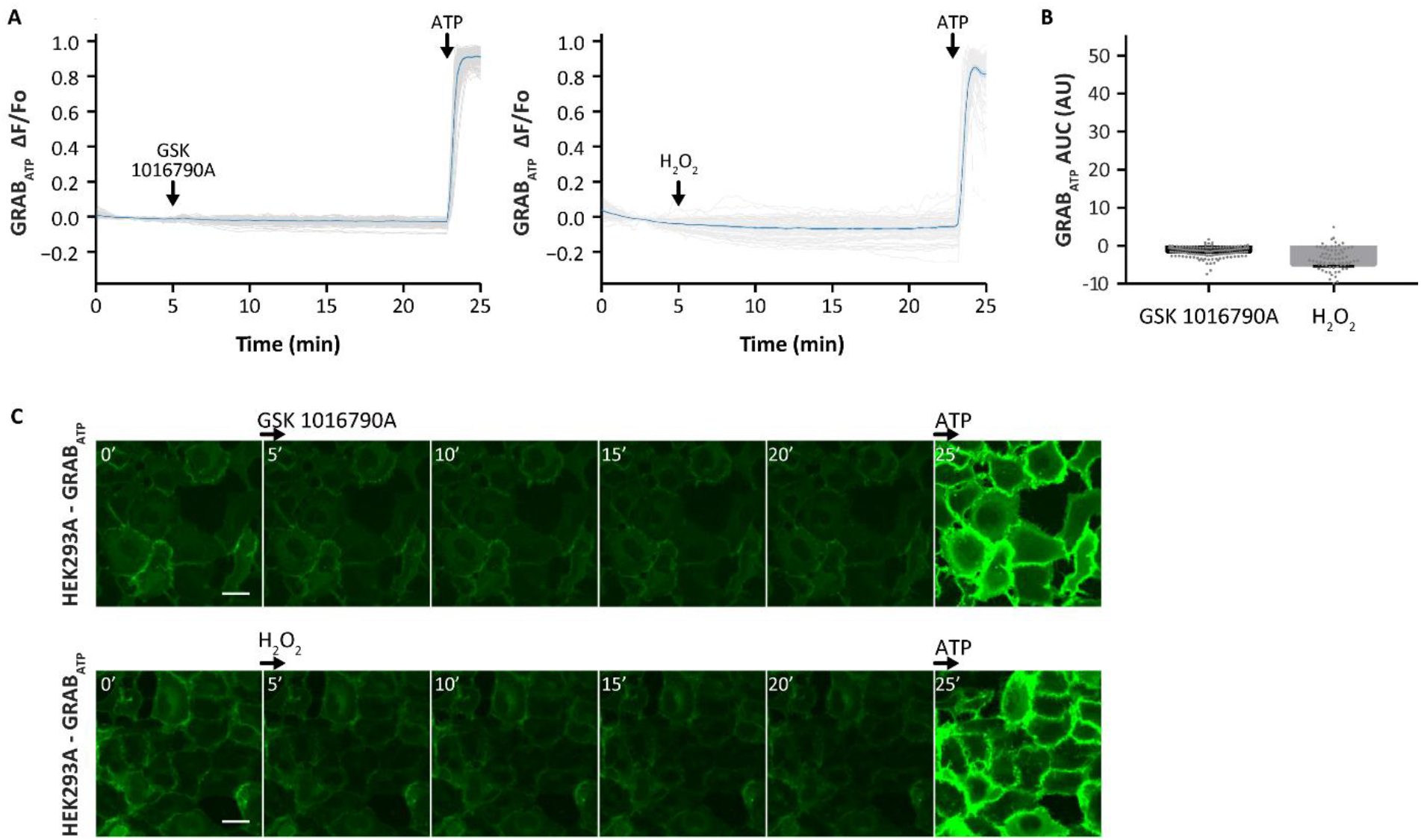
Neither GSK 1016790A, nor H2O2 evoke ATP secretion from GRABATP sensor expressing HEK293A cells. (A) After a wash to remove any residual ATP in the media, the cells were treated either with 2.5 nM GSK 1016790A or 500 μM H2O2, and then with 1 μM ATP, as a positive control. Gray lines represent the normalized mean GRABATP intensity of every cell over time. Blue line shows the overall average of the normalized mean intensity of all the cells ± SEM (GSK 1016790A: n = 149, H2O2: n = 83 cells from 3 independent experiments). (B) The area under the curve (AUC) was measured for each cell during the time of the GSK 1016790A and H2O2 stimulus, between 5-27 min, and was plotted per condition (mean ± SEM, gray dots represent values of individual cells). (C) Fluorescent images from different time points. Scale bars: 25 μm.

**Figure 8 – figure supplement 1.**
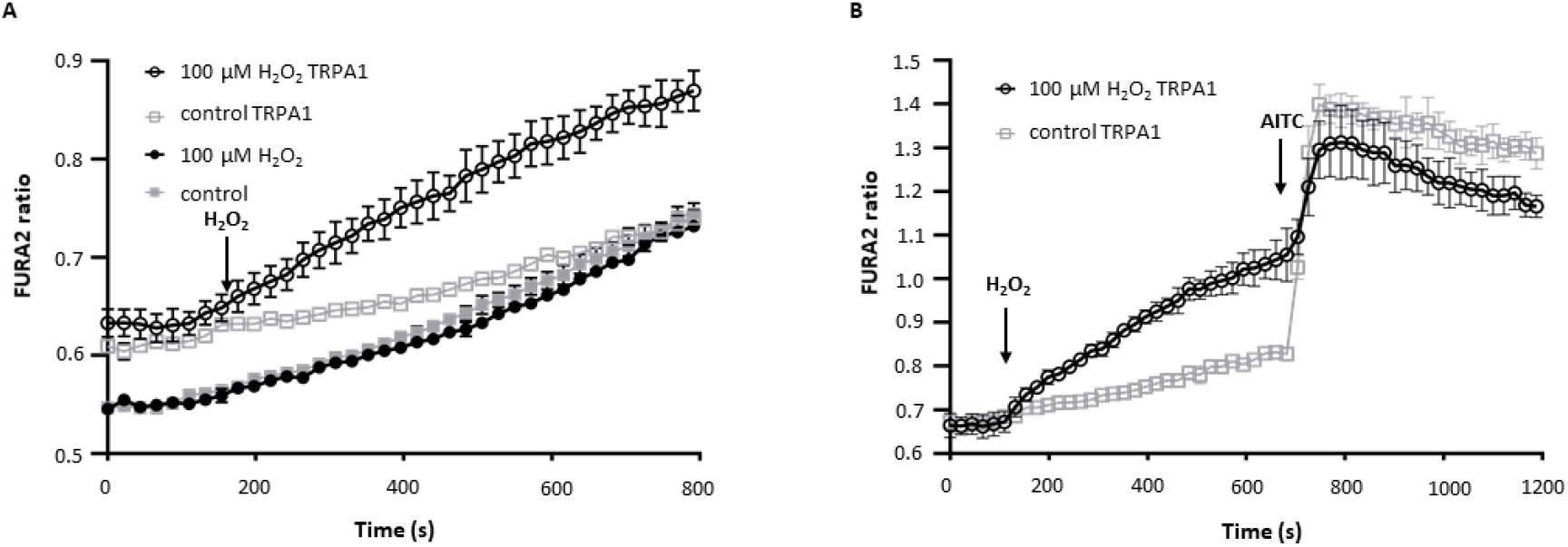
Ca^2+^ measurements with Fura-2-AM in HEK293 cells expressing TRPA1. (*A*) TRPA1-dependent effect of H_2_O_2_ on [Ca^2+^]_ic_. [Ca^2+^]_ic_ responses evoked by 100 μM H_2_O_2_ in the presence of TRPA1. (*B*) Change in [Ca^2+^]_ic_ of TRPA1 transfected HEK cells in response to sequential applications of 100 μM H_2_O_2_ and 10 μM AITC. Representative plot from at least 3 independent experiments shows mean ± SD of triplicate.

